# Polymorphic α-Glucans as Structural Scaffolds in *Cryptococcus* Cell Walls for Chitin, Capsule, and Melanin: Insights from ^13^C and ^1^H Solid-State NMR

**DOI:** 10.1101/2025.04.18.649559

**Authors:** Ankur Ankur, Jayasubba Reddy Yarava, Isha Gautam, Faith J. Scott, Frederic Mentink-Vigier, Christine Chrissian, Li Xie, Dibakar Roy, Ruth E. Stark, Tamara L. Doering, Ping Wang, Tuo Wang

## Abstract

*Cryptococcus* species are major fungal pathogens responsible for life-threatening infections in approximately a million individuals globally each year, with alarmingly high mortality rates. These fungi are distinguished by a distinctive cell wall architecture further reinforced by two virulence-associated layers, melanin and capsule, rendering them insensitive to antifungal agents targeting the cell wall, such as echinocandins. The molecular interplay between these three biomolecular layers remains poorly understood. Here we employ solid-state NMR spectroscopy to examine intact cells of both wild-type and capsule-deficient strains of *C. neoformans*, along with its melanized cells. High-resolution ^13^C and ^1^H data revealed five distinct structural forms of α-1,3-glucans that play versatile roles in forming the rigid cell wall scaffold by interacting with chitin microfibrils and chitosan, and in stabilizing the mobile matrix by associating with β-1,6-glucan and a small fraction of β-1,3-glucan. Two primary forms of α-1,3-glucans were distributed throughout the cell wall, hosting melanin deposition in the inner domain and capsule attachment on the cell surface. These findings offer a paradigm shift in understanding the cryptococcal cell wall and its interaction with two key virulence factors on opposite sides, raising critical biochemical questions that could inform the development of more effective antifungal treatments for cryptococcosis.

## INTRODUCTION

*Cryptococcus* species are a group of encapsulated basidiomycetous fungi causing life-threatening infections in immunocompromised and immunocompetent individuals^1^. These fungi produce characteristic virulence factors, including the antiphagocytic polysaccharide capsule, antioxidant melanin pigment, and extracellular proteases, including ureases and phospholipases^2,3^. Each year, approximately 200,000 cases of cryptococcal meningitis are reported worldwide, with mortality rates of 20-70% that vary significantly depending on disease severity and patient health status^4,5^. Untreated diseases are uniformly fatal. *Cryptococcus neoformans* and *Cryptococcus gattii* are the primary species responsible for these infections, with *C. neoformans* accounting for 90% of clinical isolates^6,7^. Pulmonary infections are established upon inhalation of spores or desiccated yeast cells^8^. In immunocompromised individuals, particularly when associated with HIV/AIDS, the infection can disseminate, enabling the pathogen to traverse the blood-brain barrier and induce meningoencephalitis—an inflammatory condition affecting the brain and its surrounding tissues^9,10^.

The standard treatment regimen for cryptococcosis involves amphotericin B, flucytosine, and fluconazole, typically administered for 6 to 12 months; however, this prolonged therapy is both financially burdensome and associated with significant adverse effects^11,12^. Recent antifungal research has focused on targeting fungal cell wall biosynthesis, leading to the successful development of echinocandins, which inhibit β-1,3-glucan synthesis—an essential structural component of most fungal cell walls^13^. However, echinocandins exhibit limited efficacy against *Cryptococcus* species, with the underlying mechanisms for their reduced activity remaining poorly understood but likely attributed to the unique cell wall architecture of *Cryptococcus* species^14,15^.

The cryptococcal cell wall is a dynamic, multilayered structure essential for virulence, immune evasion, and stress resistance^8,16^. Decades of chemical and imaging analyses have led to the proposition of a two-layered model of cryptococcal cell walls, in which the inner layer comprises an alkali-insoluble meshwork of β-glucans and chitin/chitosan, and the outer layer consists of an alkali-soluble fraction containing α- and β-glucans^16-18^. Unlike in other yeasts, where β-1,3-glucan is predominant, β-glucans in the *Cryptococcus* cell wall are primarily β-1,6-linked, forming covalent linkages with β-1,3-glucan, chitin, and cell wall proteins, while β-1,3-glucan is less abundant^15,19^. The cryptococcal cell wall is further associated with two additional layers of biomacromolecules that both serve as key virulence factors: a capsular layer coating the surface and melanin deposited between the cell wall polysaccharides and the plasma membrane^20-23^.

The cryptococcal capsule, which provides structural integrity and mediates host-pathogen interactions, is primarily composed of glucuronoxylomannan (GXM), with minor fractions of glucuronoxylomannogalactan (GXMGal) and mannoproteins^24-26^. These capsular polysaccharides are found both attached to the cell wall and released as exopolysaccharides^27^. Under nutrient-deficient conditions, *Cryptococcus* polymerizes external polyphenolic compounds, leading to melanin deposition in its cell wall^28,29^. These melanin granules form layered structures that protect fungi against oxidative stress, contribute to virulence, and allow small molecules, such as sugars and amino acids, to pass while restricting access by larger antifungal compounds^30,31^. Current knowledge of capsular carbohydrates is based mainly on exopolysaccharides isolated from culture supernatants, rather than those associated with the cell wall, while studies on melanin have focused on extracts called melanin ghosts, making it challenging to understand the interaction patterns of these complex and heterogeneous polymers within the cell wall^27,32^.

Recently, solid-state NMR spectroscopy has been introduced to enable high-resolution structural analysis of polysaccharides within intact fungal cells, thereby eliminating the need for cell disruption or fractionation and allowing the investigation of native physiological architectures and interactions without perturbation. Here, we use ^13^C and ^1^H solid-state NMR, enhanced by the sensitivity-boosting Dynamic Nuclear Polarization (DNP) technique^33-36^, to explore the structural polymorphism and assembly of carbohydrate polymers in intact cells of wild-type and acapsular *C. neoformans* strains. Our detailed examination of the cell wall reveals that β-1,6-glucan is the predominant component of the mobile matrix, accompanied by smaller amounts of β-1,3-glucans and mannoproteins, and is also present—albeit less abundantly—in the rigid phase alongside chitin, chitosan, and α-1,3-glucans. α-1,3-glucans dominate the rigid core and exhibit five distinct structural forms, each contributing uniquely to mechanical reinforcement, mobile matrix integration, capsular polysaccharide anchoring, and melanin deposition. These novel structural insights elucidate the organization of *C. neoformans* cell walls, provide a molecular perspective on their interface with melanin and the capsule, and underscore the critical and diverse structural roles of α-glucans, highlighting their potential as novel therapeutic targets.

## RESULTS

### Predominance of α-1,3-glucans in the rigid core of cell wall in *Cryptococcus*

*Cryptococcus* species were initially classified into subtypes based on capsular polysaccharide antigenicity, but are now distinguished by DNA sequencing, ecological characteristics, and pathobiological evidence^37-40^. In this study, we selected several representative strains of the dominant pathogen *C. neoformans* for examination. To enable studies of the capsule, we compared a clinical wild-type strain H99 with an acapsular *pka1* mutant, both of serotype A and in the same background, as well as an environmental strain JEC20 with a hypocapsular Cap70 mutant, both of serotype D (also termed *C. deneoformans*) and in the same background^41,42^. These mutants either completely lack or significantly suppress capsule formation on the cell surface, as confirmed by SEM imaging (**Fig. 1a**). Correspondingly, the mean of total cell diameter decreased from 5.7 µm in H99 cells to 4.6 µm in *pka1* cells and from 5.9 µm in JEC20 cells to 4.2 µm in Cap70 cells (**Fig. 1b** and **Supplementary Table 1**).

**Figure 1.**
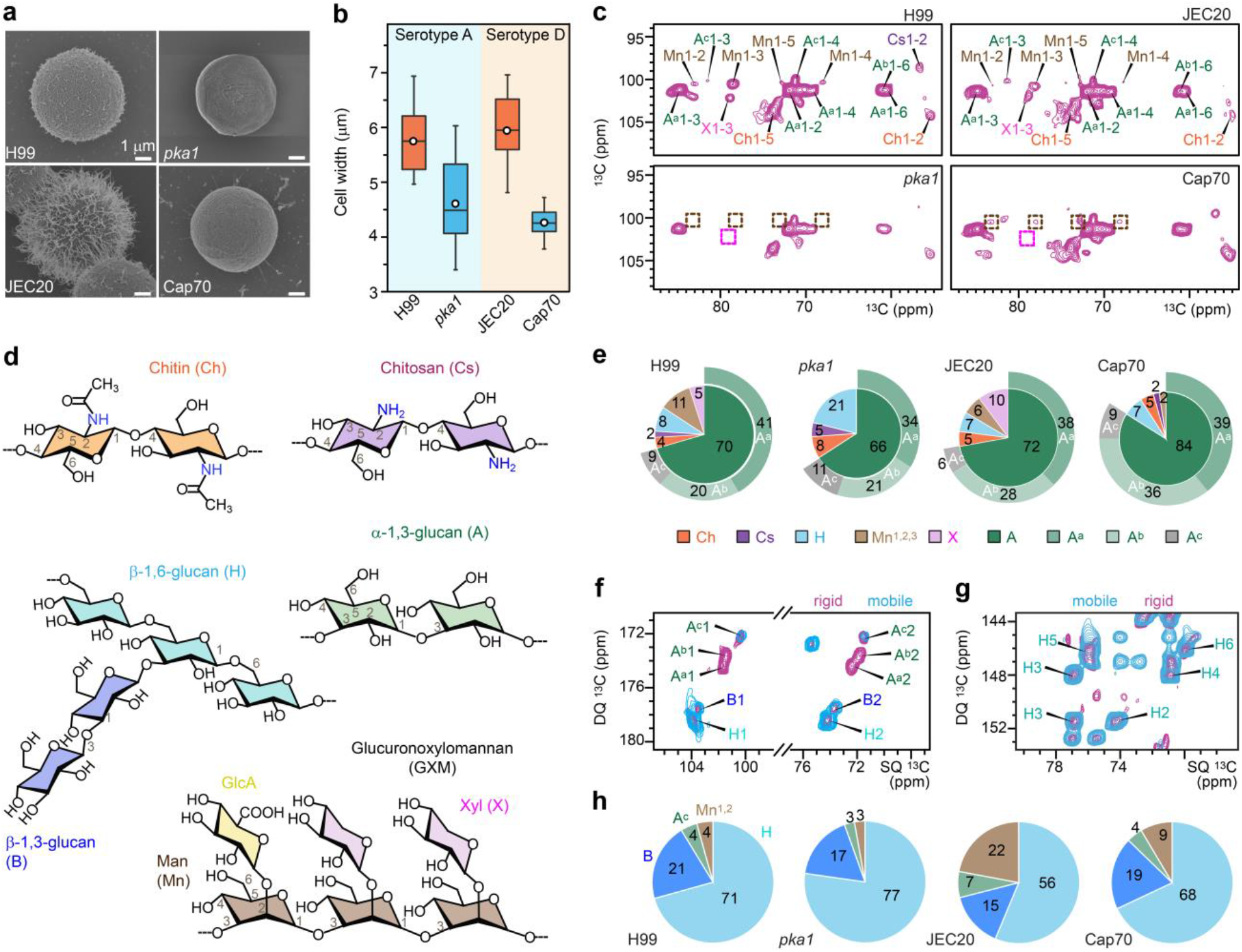
Polysaccharide composition varies in wildtype strains and capsule mutants of *C. neoformans*. (**a**) SEM images of *C. neoformans* serotype A strains (wildtype H99 and acapsular *pka1*) and serotype D (wildtype JEC20 and hypocapsular Cap70). (**b**) Total cell diameter measured from SEM images. Boxes represent the interquartile range (IQR), with whiskers extending to 1.5 times the IQR. Open circles: average; horizontal lines: median. Sample sizes: H99 and *pka1* (n = 24), JEC20 (n = 19), and Cap70 (n = 32). (**c**) 2D ^13^C-^13^C CORD correlation spectra of four *C. neoformans* samples. The absence of α-1,2,3-mannose and xylose signals in the acapsular *pka1* mutant is highlighted with dashed brown and magenta squares, respectively, with intensity reductions observed in the hypocapsular Cap70 mutant. Observed carbohydrates include α-1,3-glucan (A), chitin (Ch), chitosan (Cs), β-1,6-glucan (H), α-1,2,3-mannose (Mn), and xylose (X). Three subtypes of α-1,3-glucan (A^a^, A^b^, and A^c^) are indicated by superscripts. (**d**) Simplified structural representations of key polysaccharides, with NMR abbreviations and key carbon sites labeled. (**e**) Molar composition of rigid polysaccharides estimated from resolved cross-peak volumes in 2D ^13^C CORD spectra. (**f, g**) Overlay of refocused J-INADEQUATE spectra obtained via direct polarization (DP) for mobile molecules (cyan) and cross polarization (CP) for rigid molecules (purple) reveals (**f**) β-1,6-glucan dominance in the mobile fraction and (**g**) preferential localization of A^a^ and A^b^ in the rigid fraction, while A^c^ is distributed across both rigid and mobile phases. (**h**) Molar composition of mobile polysaccharides estimated from well-resolved peaks in DP refocused J-INADEQUATE spectra.

Rigid polysaccharides were analyzed using a 2D ^13^C-^13^C CORD correlation experiment^43^, revealing strong signals for α-1,3-glucans, along with weaker signals for chitin, chitosan, and β-1,6-glucan (**Fig. 1c, d**; **Supplementary Fig. 1** and **Table 2**). Intensity analysis indicated that α-1,3-glucans constitute 66-84% of the rigid polysaccharides across all four strains, while chitin and chitosan together account for 6-13% (**Fig. 1e** and **Supplementary Table 3**). These findings suggest that α-1,3-glucan plays a critical and unanticipated role in maintaining the mechanical scaffold of the cell wall, likely by associating with the microfibrils formed by chitin and chitosan.

Unexpectedly, signals corresponding to capsular components, including the α-1,2,3-linked mannose residues along the glucuronoxylomannan backbone and its xylose branches, were also detected within the rigid polysaccharides of H99 and JEC20 cells (**Fig. 1c** and **Supplementary Fig. 2**). These signals were entirely absent in the acapsular *pka1* mutant. In the hypocapsular Cap70 mutant, xylose signals were depleted, while α-1,2,3-linked mannose residues were retained but at reduced intensity. Previous studies have identified capsule molecules in the mobile phase of *C. neoformans* cells cultured in capsule-inducing media^44^, and our observations show that capsular polysaccharides could be structurally rigidified through interactions with cell wall polysaccharides.

### Identification of three structurally and dynamically distinct forms of α-1,3-glucans

Three magnetically distinct subtypes of α-1,3-glucans—designated as types a, b, and c—were unequivocally identified within the rigid fraction, exhibiting a sequentially decreasing prevalence within each sample (**Fig. 1c** and **Supplementary Fig. 3**). Type-a represents the most prevalent allomorphic form, comprising 34-41% of the rigid carbohydrate fraction across all samples (**Fig. 1e**). Type-b accounts for 20-36% of the rigid fraction, while type-c extends from the rigid phase into the mobile phase, constituting 6-11% of the rigid fraction (**Fig. 1e**) and 3-7% of the mobile fraction, as demonstrated later. In Cap70 cells, type-b α-1,3-glucan was remarkably enriched, reaching levels comparable to type-a in this mutant, suggesting a possible compensatory adaptation to the hypocapsular state.

The three spectroscopically distinguishable forms of α-1,3-glucans arise from local structural perturbations, including conformational variations and differential molecular interactions with neighboring components. Type-a and type-b exhibited identical ^13^C chemical shifts at C1 and C3 (101.4-101.6 ppm and 85.0-85.5 ppm, respectively), the carbon sites involved in glycosidic linkages between adjacent sugar units along the polymer chain, indicating that they share the same helical screw conformation. However, these two forms are best distinguished by their C4, C5, and C6 chemical shifts, which reflect differences in hydroxymethyl conformation related to the exocyclic -CH_2_OH group. In contrast, type-c exhibited significantly lower ^13^C chemical shifts at C1 (100.4 ppm) and C3 (81.9 ppm), indicating an entirely restructured helical screw conformation compared to the other forms.

These α-1,3-glucan allomorphic forms also have distinct dynamic properties: while all three were identified within the rigid fraction, a small amount of type-c was also detected in the mobile fraction (**Fig. 1f** and **Supplementary Figs. 3, 4**). This specific form of α-glucan may serve as a transitional component between the rigid α-glucan and chitin domains and the dynamic matrix primarily composed of β-1,6-glucan (56-77% of the mobile fraction; **Fig. 1g, h**), β-1,3-glucan (15-21%), and some α-1,2-linked mannose residues, likely derived from mannoproteins (**Fig. 1h** and **Supplementary Table 4**). This finding provides evidence for the critical role of α-1,3-glucan in mediating physical interactions that bridge rigid and dynamic domains.

In the mobile phase, we observed abundant β-1,6-glucan, which is expected as it is unusually rich in *Cryptococcus* compared to other yeasts such as *Saccharomyces cerevisiae*^15,19^. The structure of cryptococcal β-1,6-glucan consists of short chains, often branched with β-1,3-glucan, and our data further identified it as the dominant component of the mobile matrix. The mobile phase also contains proteins and lipids; however, their widespread distribution throughout the cell precluded a focused analysis of their specific contributions to cell wall structure (**Supplementary Fig. 5**).

### Rigid α-glucan forms interact with capsules to form dehydrated domains on cell surface

Hydration profiles of rigid biopolymers were analyzed using water ^1^H polarization transfer to carbohydrates via water-editing experiments, where the S/S_0_ intensity ratios between water-edited (S) and control (S_0_) spectra reflect the extent of water retention at specific carbon sites (**Fig. 2a** and **Supplementary Fig. 6**)^45-47^. Two α-1,3-glucan subtypes, a and b, were the least hydrated molecules in *Cryptococcus* cell walls, with average S/S_0_ values of 0.26-0.43 across all four strains (**Fig. 2a** and **Supplementary Table 5**). Meanwhile, in the examination of the molecular motions of rigid polysaccharides through NMR relaxation measurements, the resolvable carbon sites of type-a α-1,3-glucan exhibited the longest ^13^C-T_1_ and ^1^H-T_1ρ_ relaxation time constants across the four strains, with average values of 2.7-3.9 s (**Fig. 2b**; **Supplementary Fig. 7** and **Table 6**) and 9.7-12.6 ms (**Fig. 2c** and **Supplementary Fig. 8**), respectively. These values are significantly higher than those of the chitin and β-1,6-glucans in the rigid phase, indicating that α-1,3-glucans experience highly restricted molecular motion on both nanosecond and microsecond timescales. The combination of restricted dynamics and limited hydration suggests that types-a and b α-1,3-glucans aggregate into large complexes that effectively exclude bulk water. In contrast, type-c α-1,3-glucan was not only more hydrated but also more heterogeneously hydrated compared to other subforms (**Fig. 2a**). This further supports the notion that type-c α-1,3-glucan plays a crucial role in bridging the rigid and mobile phases of the α-1,3-glucan matrix (**Fig. 1e, h**).

**Figure 2.**
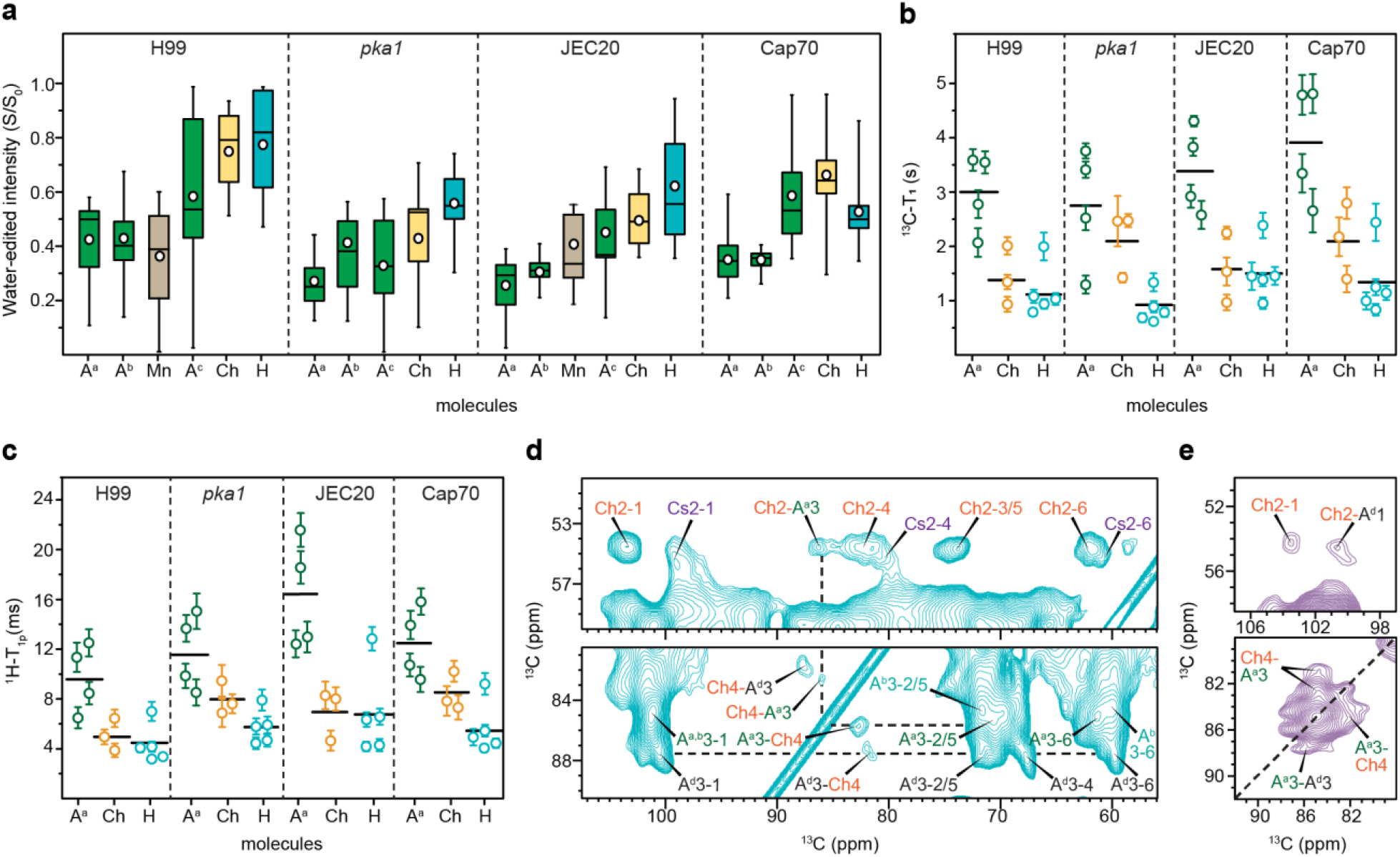
Hydration, dynamics, and DNP spectra reveal α-glucan interactions with capsule and chitin. **(a)** Intensity ratios (S/S_0_) between water-edited peak spectra (S) and control spectra (S_0_) showing the extent of water association for α-1,3-glucan (A), chitin (Ch), 1,2,3-linked mannan (Mn), and β-1,6-glucan (H), with the superscripts indicating the respective subtypes. Open circles: average; horizontal lines: median. Data size: A^a^ (n=16) in all four samples; A^b^, A^c^, Ch, H (n=9 each) in all; Mn (n=9) in H99 and JEC20 only. **(b)** ^13^C-T_1_ relaxation time constants of rigid cell wall polysaccharides. (**c**) ^1^H-T_1ρ_ relaxation time constants of rigid polysaccharides. For both panels b and c, error bars are s.d. of the fit parameters and horizontal lines represent the average. (**d**) DNP-enhanced 2D ^13^C-^13^C DARR spectrum of H99 cells. A new, minor form of α-1,3-glucan (A^d^) is resolved. Intermolecular cross peaks were observed between chitin and type-a and type-d α-1,3-glucans, with dashed lines to mark the intramolecular and intermolecular cross peaks of these two α-1,3-glucan subtypes. (**e**) DNP-enhanced 2D ^13^C-^13^C PAR spectrum with 15 ms recoupling time. Top: observation of a cross peak between chitin and A^d^. Bottom: the dashed line indicates the diagonal of the C3 sites, with off-diagonal intensities showing intermolecular cross peaks.

β-1,6-glucans are the best-hydrated rigid polysaccharide in *C. neoformans*, with high S/S_0_ averages of 0.53-0.77 (**Fig. 2a**). β-1,6-glucan also exhibited short ^13^C-T_1_ (0.9-1.5 s) and ^1^H-T_1ρ_ (4.4-6.8 ms) relaxation times— about half to two-thirds those of type-a α-1,3-glucans (**Fig. 2b, c**). β-1,6-glucan is the most dynamic molecule, maintaining substantial hydration even within the rigid core. Chitin displays intermediate rigidity and hydration profiles, typically between β-1,6-glucan and type-a α-1,3-glucan. Chitin forms a microfibrillar structure primarily through antiparallel chain packing but is not the most rigid molecule in the cell wall. This may be due to a high level of deacetylation to chitosan, estimated to be 30-40% based on molar composition (**Fig. 1e**), which confers dynamics on these microfibrils.

Unexpectedly, 1,3-linked mannose residues, which compose the backbone of GXM, exhibited low S/S_0_ values, averaging 0.38 and 0.41 in H99 and JEC20 cells, respectively. This indicates significant dehydration of capsular polysaccharides in both wild-type serotypes A and D strains (**Fig. 2a**), which is counterintuitive because capsular GXM is surface-exposed and would therefore be expected to be well-hydrated. A plausible explanation is that GXM associates with α-1,3-glucan subtypes a and b, which not only rigidifies GXM but also forms a dehydrated layer on the cell surface. This structural organization contrasts with cryptococcal capsules produced in capsule-inducing media, where GXM is loosely associated with cell wall molecules, thus remaining mobile and solubilized^44^.

Interestingly, the two serotypes responded differently to the absence or reduction of capsular polysaccharides. In serotype A, water association decreased substantially in the *pka1* mutant compared to wild-type H99 cells (**Fig. 2a**), with average S/S_0_ ratios dropping from 0.42 to 0.27 for type-a α-glucan, 0.58 to 0.33 for type-c α-glucan, 0.77 to 0.56 for β-1,6-glucan, and 0.75 to 0.43 for chitin. In contrast, serotype D exhibited increased hydration for most cell wall polysaccharides, with S/S_0_ values rising from 0.26 in JEC20 to 0.37 in Cap70 cells for type-a α-glucan, 0.45 to 0.59 for type-c α-glucan, and 0.49 to 0.67 for chitin, except for β-1,6-glucan, whose hydration level decreased. Therefore, different serotypes may exhibit variations in cell wall organization and adopt distinct compensatory mechanisms to remodel their polysaccharides in response to capsule depletion. However, a consistent observation is that chitin dynamics became more confined in both acapsular and hypocapsular mutants (**Fig. 2b, c**), suggesting strengthened interactions with α-glucans, likely due to increased availability of interaction sites on types-a and b α-glucans after GXM removal.

### Two specific α-glucans interact with chitin to form partially ordered domains in the cell wall

We used DNP to enhance the NMR sensitivity for polysaccharides in H99 cells by 11-fold through polarization transfer from electrons in the biradical AsymPol-POK to the ^1^H and then ^13^C nuclei in these cellular biomolecules^33,48^. The observed signals arose from partially ordered molecules, including chitin, chitosan, types-a and b α-glucans, and a newly identified minor form, type-d α-1,3-glucan (labeled as A^d^ in **Fig. 2d**). This new form of α-1,3-glucan exhibited unique C3 and C4 chemical shifts at 87.6 ppm and 67.8 ppm, with weak signals detectable only through DNP enhancement. Meanwhile, signals from mobile and semi-mobile molecules, such as β-1,6-glucan and type-c α-1,3-glucan, were broadened out due to a diverse ensemble of conformations trapped at the cryogenic temperature.

With the enhanced sensitivity, strong intermolecular interactions were detected between chitin and types-a and type-d α-1,3-glucans, as shown by the unambiguous Ch2-A^a^3, Ch4-A^a^3, A^a^3-Ch4, Ch4-A^d^3, and A^d^3-Ch4 cross-peaks observed in the 100-ms dipolar-assisted rotational resonance (DARR) spectrum (**Fig. 2d**). We also observed three cross peaks in a 15-ms proton-assisted recoupling (PAR) spectrum, including Ch2-A^d^1, Ch4-A^a^3, and A^a^3-Ch4, further confirming the interactions between chitin and these two specific forms of α-1,3-glucans (**Fig. 2e**)^49,50^. In addition, a cross peak between the carbon-3 sites of type-a and type-d α-1,3-glucans (A^a^3-A^d^3) was detected (**Fig. 2e**). Together, these observations confirm that type-d and type-a α-1,3-glucans are associated with each other and further packed with chitin in the rigid domain of the cell wall, while type-b and type-c α-1,3-glucans are not colocalized with chitin microfibrils.

### Two structural forms of α-1,3-glucans host melanin deposition

In addition to the capsule, another crucial virulence factor of *Cryptococcus* species is their ability to synthesize melanin, which is deposited in the cell wall, and forms a protective barrier against environmental and host stressors^31^. Melanin polymers are proposed to be composed of stacked aromatic and indolic rings, formed via the oxidative polymerization of L-3,4-dihydroxyphenylalanine (L-DOPA) catalyzed by the laccase enzyme (**Fig. 3a, b**)^51,52^. Since *Cryptococcus* requires an exogenous substrate for melanin biosynthesis, we supplemented the minimal medium with 1 mM ring-^13^C_6_-labeled L-DOPA to cultivate melanin-rich *C. neoformans* H99 cells^53-55^. Consequently, the melanin-labeled cells exhibited enhanced aromatic ^13^C signals in the 110-160 ppm range, particularly in the 140-150 ppm region, corresponding to non-protonated melanin carbon sites (**Fig. 3c**, see expanded spectral region). These distinct melanin aromatic signals enabled us to establish correlations with carbohydrate proton resonances to map out melanin-carbohydrate interactions within intact cells.

**Figure 3.**
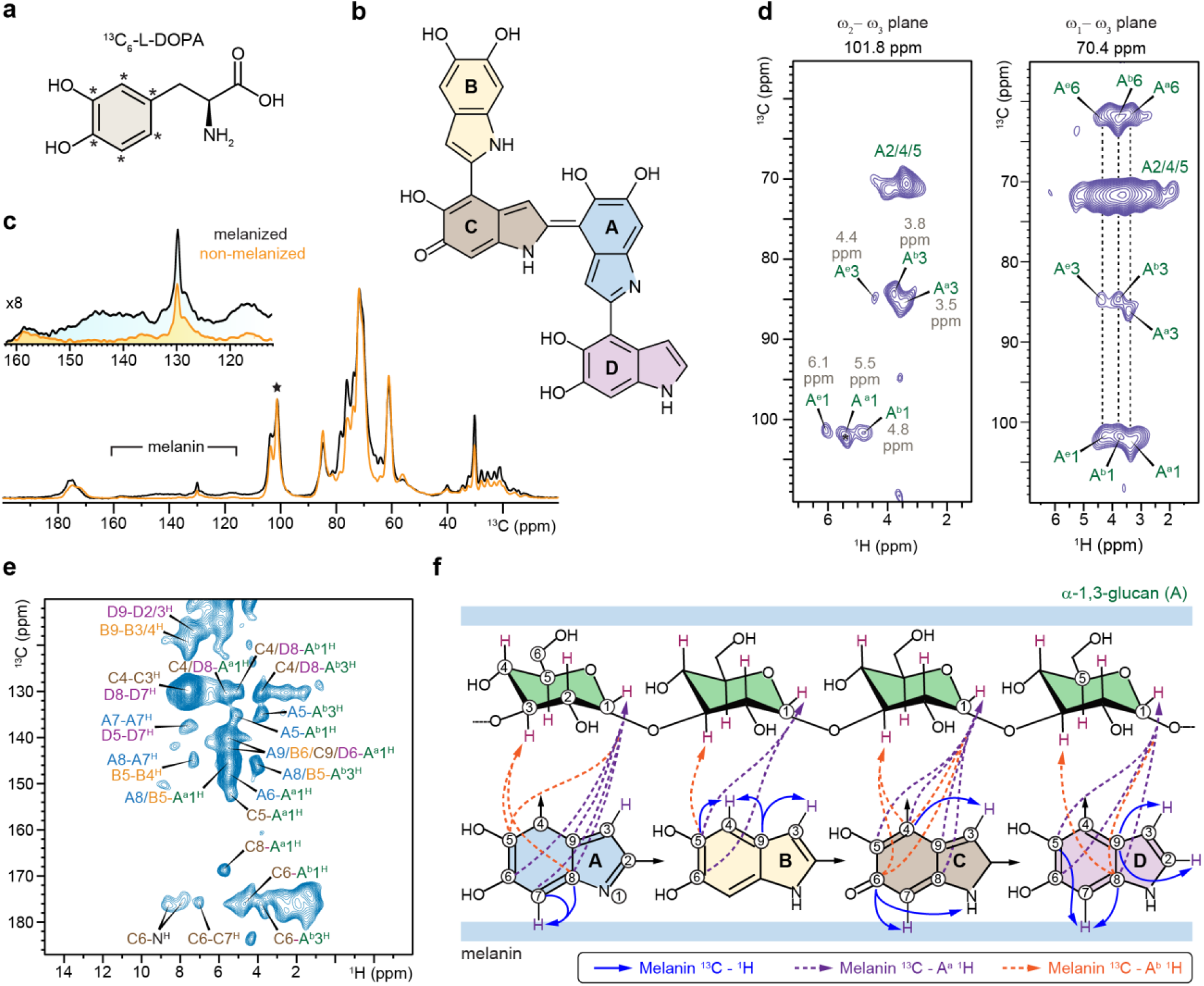
^1^H-detected solid-state NMR shows interactions of α-1,3-glucan with melanin. (**a**) Structure of melanin precursor ring-^13^C_6_-labeled L-DOPA. Asterisks indicate the ^13^C-labeling sites. (**b**) Representative structure of DHI melanin formed by covalently linked subunits. (**c**) 1D ^13^C CP spectra of non-melanized (orange) and melanized (black) *C. neoformans* H99 cells. The spectra are normalized by the α-1,3-glucan carbon 1 peak at 101 ppm (asterisk). The zoom-in region shows the aromatic signals from melanin, with intensities magnified by 8-fold. (**d**) Polymorphic forms of α-1,3-glucans (A^a^, A^b^, and A^e^) resolved using ^1^H-detected 3D hCCH TOCSY (WALTZ-16) spectrum. ^1^H chemical shifts (ppm) are in grey. Key strips are extracted for ω_2_-ω_3_ (^13^C-^1^H) and ω_1_-ω_3_ (^13^C-^1^H) planes. (**e**) 2D hChH with 0.8 ms RFDR mixing showing intermolecular interaction between melanin non-protonated carbons and α-1,3-glucan protons. (**f**) Structural summary of observed interactions between cell wall α-1,3-glucan and melanin units. Carbon and nitrogen atom numbering in melanin fragments follows the standard convention for indoles. Blue solid lines: cross peaks observed between non-protonated carbons and protons in protonated sites within melanin. Dashed lines in purple and orange indicate the intermolecular interactions between melanin carbons and protons in type-a and type-b α-1,3-glucans, respectively. The experiments were performed on 600 MHz NMR at 60 kHz MAS.

We further identified three polymorphic forms of α-1,3-glucan in melanized *C. neoformans* cells, designated A^a^, A^b^, and A^e^, from the ^13^C-^1^H strips extracted from the 3D hCCH TOCSY (WALTZ-16) spectrum (**Fig. 3d** and **Supplementary Table 7**). These polymorphs exhibited distinct ^1^H chemical shifts: the anomeric carbon (A1) correlated with three distinct ^1^H resonances at 5.5 ppm (A^a^), 4.8 ppm (A^b^), and 6.1 ppm (A^e^) in the ω_2_-ω_3 13_C-^1^H strip, while the C3 position correlated with protons at 3.5 ppm (A^a^), 3.8 ppm (A^b^), and 4.4 ppm (A^e^) (**Fig. 3d**). The ω_1_-ω_3 1_H-^13^C strip extracted from 70.4 ppm further revealed the through-bond carbon connectivity of these three polymorphic forms and established correlations with distinct carbon sites, such as the resolvable carbons at positions 3 and 1 (**Fig. 3d**). In this analysis, the A^c^ and A^d^ forms detected in ^13^C-based experiments were not observed in this analysis due to their semi-dynamic nature or low abundance.

Next, we measured a 2D hChH spectrum, which employed an extended, 0.8 ms radio frequency driven recoupling (RFDR) mixing period to facilitate long-range ^1^H-^1^H polarization transfer (**Fig. 3e**). Notably, this experiment facilitated the observation of extensive correlations between non-protonated carbons and protons at protonated sites within the melanin structure (**Fig. 3e**; blue solid lines in **Fig. 3f**). The ^13^C and ^1^H chemical shifts identified in melanin are summarized in **Supplementary Fig. 9** and **Tables 8** and **9**. In melanin subunit D, key cross-peaks include those between carbon-9 and protons-2/3 (D9-D2/3^H^), as well as D5-D7^H^ and D8-D7^H^ (**Fig. 3e, f**). Similar cross-peaks were observed within other melanin subunits, such as A7/8-A7^H^, B9-B3/4^H^, B5-B4^H^, C4-C3^H^, and C6-C7^H^. The C6-N^H^ cross-peak further revealed spatial proximity between C-fragment carbons and indole amide protons from the same or neighboring subunits.

Importantly, the hChH experiment simultaneously revealed extensive intermolecular interactions between the carbons in the indole rings of melanin and the protons in carbohydrates. For example, the carbon 4 of melanin subunit-C and carbon 8 of subunit-D, two non-protonated carbons resonating at 130 ppm, exhibited correlations with the proton 1 of type-a α-1,3-glucan at 5.5 ppm, the proton 1 of type-b α-1,3-glucan at 4.8 ppm, and the proton 3 of type-b α-1,3-glucan at 3.8 ppm, resulting in the C4/D8-A^a^1^H^, C4/D8-A^b^1^H^, and C4/D8-A^b^3^H^ cross peaks (**Fig. 3e**). Type-a and type-b α-1,3-glucans showed 13 and 10 strong cross-peaks with melanin, respectively, including A8/B5-A^a^1^H^, A5-A^b^3^H^, A5-A^b^1^H^, A6-A^b^1^H^, C5-A^a^1^H^, C8-A^a^1^H^, C6-A^b^1^H^, and C6-A^b^3^H^, in addition to those previously described (**Fig. 3e, f**). These intermolecular interactions provide the first molecular-level experimental evidence that α-1,3-glucans act as a previously unrecognized partner with melanin, with their type-a and type-b structural variants serving as the primary interactors.

## DISCUSSION

Recent advances in solid-state NMR spectroscopy have significantly enhanced our ability to investigate fungal cell walls, offering detailed insights into the structures of wall polymers, as well as their hydration, dynamics, and intermolecular interactions^56-60^. Such analysis describes molecular behavior in the context of intact cells, in a way that prior methods, which relied on physical, chemical, and/or enzymatic perturbation, could not^61,62^. The application of these techniques to the major fungal pathogen *C. neoformans* in this study has uncovered unexpected features of the cell wall and revealed its interactions with two critical virulence factors: the extracellular capsule and cell-associated melanin.

Previous studies of cryptococcal cell walls, based on structural and imaging analyses, suggested a bilayer structure with an inner layer composed of interlinked chitin, chitosan, and β-glucans and an outer layer composed of α- and β-glucans^16,17,63^. In this model, melanin localizes to the inner aspect of the wall, near the plasma membrane, while the capsule polysaccharide is associated with α-1,3-glucan in the outer layer^23,25^. The integration of our NMR data with this existing model now supports major revisions (**Fig. 4**). For example, our data demonstrate that α-1,3-glucan is a key structural component of cryptococcal cell walls, occurring both in the inner wall, where it interacts with melanin and chitin, and in the outer wall, where it promotes capsule attachment. We also found that β-1,6-glucan, an under-investigated component in the fungal cell wall^64^, is distributed across dynamically distinct domains. It is primarily localized in the mobile phase, where it forms the main component of the soft matrix, with smaller fractions extending into the rigid phase to contribute to structural integrity. This also explains the limited efficacy of echinocandins against Cryptococcus species, as β-1,3-glucan is present in low abundance, with β-1,6- and α-glucans dominating, and β-1,3-glucan being confined exclusively to the mobile phase.

**Figure 4.**
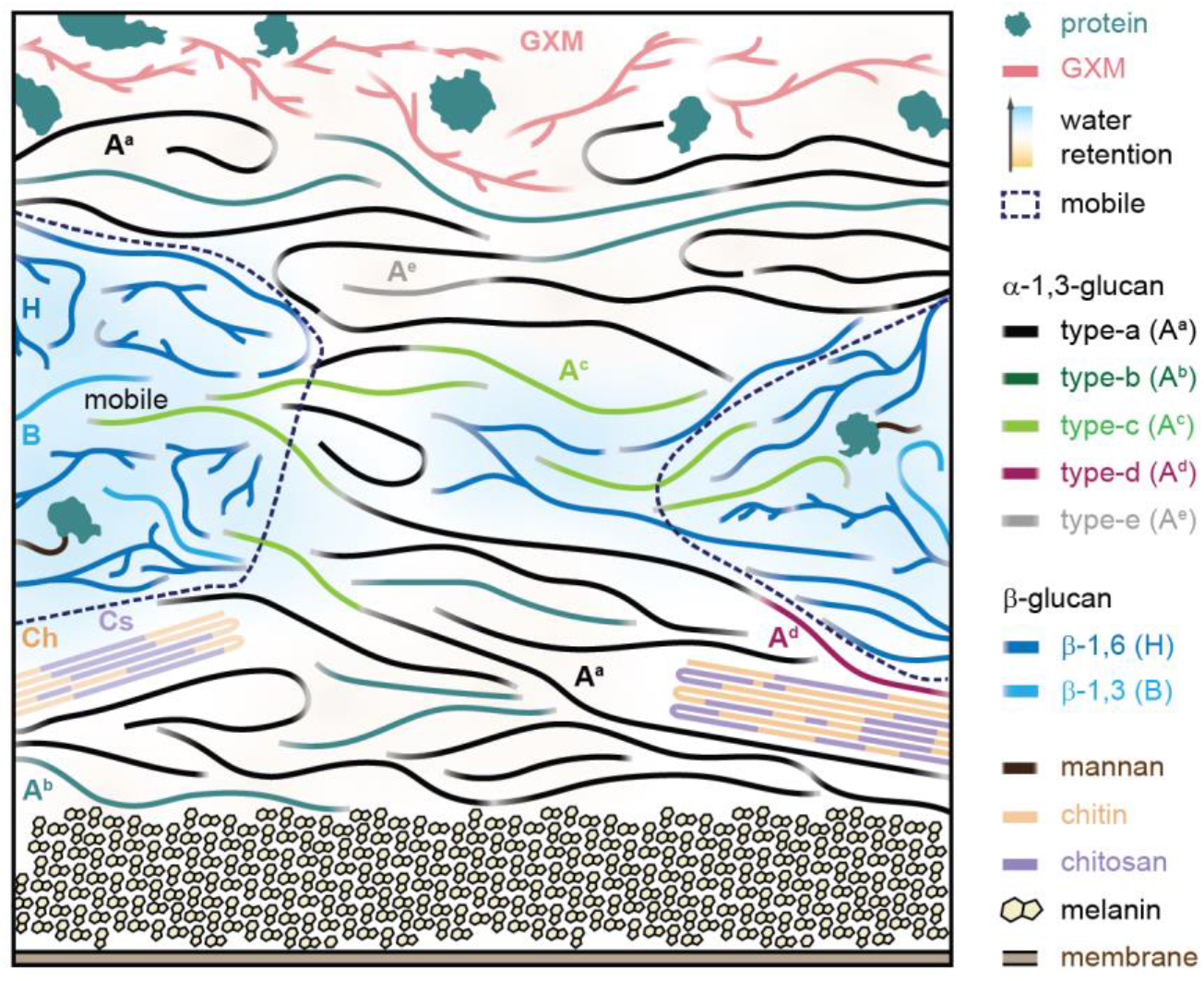
Organization of cryptococcal cell walls and association with capsule and melanin. The illustrative scheme is based on NMR data from H99 cells, integrated with previously reported biochemical and imaging analyses^16,31,63^. Five distinct forms of α-1,3-glucans constitute 70% of the rigid polysaccharides. Capsular polysaccharides, primarily GXM, are associated with type-a and type-b α-1,3-glucans in the cell wall. Melanin is stabilized by type-a and type-b α-1,3-glucans. Molecules associated with type-a and type-b α-1,3-glucans are rigidified and dehydrated. Chitin and chitosan, which together account for only 6% of rigid molecules, exhibit a high deacetylation level of approximately 30% and are stabilized through interactions with type-d and type-a α-1,3-glucans, both of which colocalize. The mobile phase (highlighted by dashed lines) is predominantly composed of β-1,6-glucan (70%), which regulates the water association of cryptococcal cell wall, with a minor fraction of β-1,3-glucan, present either as linear chains or as sidechains of β-1,6-glucan. Additionally, type-c α-1,3-glucan extends through rigid and mobile phases. Type-e α-1,3-glucan represents a minor structural variant that does not exhibit an association with melanin. Molecular fractions are considered in this model, though the representation is not strictly to scale. The background gradient illustrates the water distribution within the cell wall.

Notably, the cell wall α-1,3-glucan of *C. neoformans* occurs in five distinct morphotypes, which we were able to resolve using high-resolution ^13^C solid-state NMR, combined with advanced ^1^H detection and DNP sensitivity enhancement (**Fig. 4**). Type-a and type-b α-1,3-glucans are the predominant structured forms, comprising 55-75% of all rigid polysaccharides (**Fig. 1e**). These forms aggregate into partially dehydrated bundles, facilitating capsule deposition on the cell surface and rigidifying the capsule polysaccharide GXM (**Fig. 2a-c**; upper portion of **Fig. 4**). This highlights the essential role of α-1,3-glucan in capsule-cell wall attachment, and provides a structural explanation for why disruption of the α-1,3-glucan synthase gene results in viable *C. neoformans* cells lacking a surface capsule, despite the successful production of capsule components^65^.

Surprisingly, these major forms of α-1,3-glucan (types a and b) that interact with capsule polysaccharide also serve to host melanin deposition (**Fig. 3e**; bottom portion of **Fig. 4**). These unexpected findings lead to two novel conclusions. First, they solve the mystery of which wall component helps organize melanin deposition, which was not known despite the implication of the cell wall in this process^29,51^. Second, they indicate that these two forms of α-1,3-glucans exhibit a broad spatial distribution, localized both at the cell surface, where they mediate capsule association, and near the plasma membrane, where they colocalize with melanin (**Fig. 4**). This bimodal distribution significantly shifts the structural paradigm of cryptococcal cell walls, revealing that the inner layer, previously believed to consist of chitin, chitosan and β-glucan, contains additional components^16^. This model is also consistent with the persistent melanin-cell-wall association observed even in samples with relatively low chitin and chitosan content.

The remaining three forms of α-1,3-glucan, though less abundant, are also functionally significant. Type-c is present in both the rigid and mobile phases (**Fig. 1e, h**), and thus plays a critical role in integrating α-1,3-glucan aggregates into the mobile matrix. Type-d α-1,3-glucan is a minor but highly ordered form that colocalizes with type-a α-glucan, both of which are physically associated with chitin microfibrils (**Fig. 2d, e**). Type-e, another minor form, underlies the ^13^C signals of type-a and type-b α-glucans and is distinguishable by ^1^H chemical shifts, indicating local structural variations compared to the predominant forms (**Fig. 3d**).

Our results provide an unprecedented understanding of the structural complexity of α-1,3-glucan in the construction of cryptococcal cell walls and highlight its association with two key virulence factors anchored at opposite aspects of the cell wall. The multiple roles of this versatile polymer may also explain its unusually high abundance in *Cryptococcus* species. Meanwhile, these findings raise new questions. It remains unclear whether the observed structural forms result from complexity in α-1,3-glucan biosynthesis and whether they are universally present across different *Cryptococcus* species, other serotypes, and other pathogenic fungal species. It also remains to be determined whether inhibitors of α-1,3-glucan synthases or other enzymes involved in biosynthesis and modification could be developed as therapeutic targets. Addressing these questions would advance our understanding of cryptococcal cell wall biosynthesis and structure, providing molecular insights for the development of effective antifungal therapies against cryptococcosis.

## METHODS

### Preparation of _13_C,_15_N-labeled fungal material

*C. neoformans* strains H99, JEC20, *pka1*, and Cap70 were cultured in ^13^C, ^15^N-enriched growth media. The growth medium consisted of 1.7% Yeast Nitrogen Base (YNB), 2.1% ^13^C-glucose (Cambridge Isotope Laboratories, CLM-1396-PK), and 1.5% ^15^N-ammonium sulfate (Cambridge Isotope Laboratories, NLM-713-PK), with the pH adjusted to 5.5 using 1M HEPES buffer. Cells were cultured in 100 mL liquid medium in 250-mL Erlenmeyer flasks and incubated at 37°C with shaking at 150 rpm (1364 × g). Fungal biomass was harvested by centrifugation at 4,000 rpm (13,700 × g) for 5 min at 4°C and the pellet was washed four times using nano-purified water followed by the centrifugation procedures. For solid-state NMR characterization, 35-45 mg of natively hydrated whole-cell material was packed into a 3.2-mm magic-angle spinning (MAS) rotor (Cortecnet, HZ16916).

To generate melanin-rich *C. neoformans* cells, H99 cells was incubated for 10-14 days at 30°C with shaking at 150 rpm in a minimal medium (29.4 mM KH_2_PO_4_, 10 mM MgSO_4_, 13 mM glycine, 3 µM thiamine, 15 mM ^13^C-glucose, pH 5.5) supplemented with 1 mM ^13^C_6_-L-DOPA (Cambridge Isotope Laboratories, CLM-1007-PK)^53,54^. Melanized fungal cells were collected and washed following the protocol described above and packed into 3.2 mm and 1.3 mm MAS rotors (Cortecnet, HZ14752) for solid-state NMR characterization.

### Scanning electron microscopy

A small amount of the same *C. neformans* material used for NMR analysis were suspended in distilled water, mixed with 4% glutaraldehyde in 0.1 M sodium phosphate buffer (pH 7.4), and fixed for 30 min at 4°C. To prepare coverslip-mounted samples, 1% Poly-L-Lysine (Sigma Aldrich, P1399) was applied as a single drop onto a plastic Petri dish, over which a 12 mm round glass coverslip was placed and allowed to adhere for 10 min. The coverslip was then removed, gently rinsed with water, and drained while preventing complete drying. A drop of the fixed cell suspension was placed on the previously poly-lysine-exposed surface of the coverslip and allowed to settle for 10 minutes. The sample was subsequently rinsed with water and dehydrated through a graded ethanol series (25%, 50%, 75%, 95%) for 10 min at each step, followed by three 10-min changes in 100% ethanol and critical point drying using a Leica Microsystems EM CPD300 critical point dryer (Leica Microsystems, Vienna, Austria) with carbon dioxide as the transitional fluid. Dried samples were mounted on aluminum stubs, sputter-coated with osmium, and examined using a JEOL 7500F field emission scanning electron microscope operating at 15 kV. Total cell diameter was quantified using ImageJ software (version V1.8.0_172) and reported in **Supplementary Table 1**.

### ^13^C solid-state NMR experiments

High-resolution 1D and 2D solid-state NMR experiments were conducted at 800 MHz (18.8 Tesla) at Michigan State University. All ^13^C-detection experiments were performed using a 3.2 mm HCN probe at 15 kHz MAS, with ambient temperatures between 283 K and 298 K. ^13^C chemical shifts were externally referenced by calibrating the adamantane CH_2_ peak to 38.48 ppm, and the resulting spectral reference (sr) value was applied to fungal spectra. Unless otherwise specified, typical radiofrequency field strengths were 83-100 kHz for ^1^H decoupling, 62.5 kHz for ^1^H hard pulses, and 50-62.5 kHz for ^13^C. Experimental parameters for all NMR spectra are documented in **Supplementary Table 10**.

1D ^13^C spectra were acquired using various polarization methods to probe molecular dynamics. Rigid components were detected via dipolar-based ^13^C cross-polarization (CP) with a 1-ms contact time, adhering to Hartmann-Hahn match conditions of 62.5 kHz for ^13^C and ^1^H. For quantitative analysis, 1D ^13^C direct polarization (DP) experiments were conducted with a long recycle delay of 35 s, ensuring full longitudinal relaxation before the next scan. Additionally, a shorter recycle delay of 2 s in the same ^13^C DP experiment enabled selective detection of mobile molecules with fast ^13^C-T_1_ relaxation. These 1D experiments were performed on three independently prepared *C. neoformans* sample replicates, demonstrating high reproducibility (**Supplementary Fig. 10**).

To facilitate resonance assignment, 2D ^13^C-^13^C correlation experiment was conducted using a 53-ms CORD (combined R2_n_^v^-driven) mixing period, which revealed intramolecular cross-peaks between carbon sites within each molecule (**Supplementary Fig. 1**)^43^. Additionally, 2D ^13^C refocused J-INADEQUATE experiments were conducted using either CP or DP (1.5 s recycle delay) to probe molecular domains with differing mobility (**Supplementary Fig. 4**)^66^. This experiment correlated double-quantum (DQ) chemical shifts with two corresponding single-quantum (SQ) shifts, generating asymmetric spectra that enabled efficient through-bond carbon-connectivity tracking. To optimize carbohydrate detection, the τ period (out of four) was set to 2.3 ms. All 1D experiments, as well as 2D CORD experiments, were conducted for all *C. neoformans* strains. Assigned chemical shifts for rigid and mobile polysaccharides are documented in **Supplementary Table 2**. Data acquisition was conducted using Topspin 3.5, and spectral analysis was performed in Topspin 4.0.8. Figures were prepared using Adobe Illustrator CS6 (V16.0.0).

Molecular composition analysis was conducted by selecting only well-resolved signals in 2D ^13^C CORD spectra for rigid components and ^13^C DP refocused J-INADEQUATE spectra for mobile molecules (**Supplementary Tables 3** and **4**). Peak volumes were quantified using the integration function in Bruker Topspin, with quantification based on the mean of resolved signals. Relative polysaccharide abundance was determined by normalizing the sum of integrals with their respective counts^67,68^. The standard error for each polysaccharide was calculated by dividing the standard deviation of integrated peak volumes by the total cross-peak count. The overall standard error was computed as the square root of the sum of squared errors for each polysaccharide. The percentage error was determined by normalizing the standard error with the average integrated peak volume and adjusting for each polysaccharide’s relative abundance^67,68^.

### Solid-state NMR of polymer hydration and dynamics

All polymer hydration and dynamics experiments were conducted on a Bruker Avance Neo 400 MHz (9.4 T) NMR spectrometer at Michigan State University using a 3.2 mm HCN MAS Bruker probe at 280 K. Water accessibility of polysaccharides was assessed using 1D and 2D water-edited ^13^C-^13^C correlation spectra^45,47^. A ^1^H-T_2_ filter (0.6-1.2 ms, strain-dependent) was applied to suppress carbohydrate signals to less than 10% but preserve 82-88% of water magnetization. (**Supplementary Fig. 6**), followed by ^1^H-^1^H mixing to transfer water ^1^H magnetization to hydrated carbohydrates. ^13^C detection was achieved via CP with a 1-ms contact time. For 2D water-edited experiments, a 4 ms ^1^H mixing period and 50 ms DARR mixing were used. Intensity ratios (S/S_0_) between the water-edited (S) and control (S_0_) spectra were obtained to quantify water retention around each carbon site (**Supplementary Table 5**). ^13^C-T_1_ relaxation was measured using the Torchia-CP scheme^69^ with z-filter durations from 0.1 μs to 8 s. For ^13^C-detected ^1^H-T_1ρ_ relaxation, a Lee-Goldburg (LG) spinlock sequence combined with LG-CP suppressed ^1^H spin diffusion, enabling site-specific measurements via bonded ^13^C detection^70,71^. For both measurements, peak intensity decay was fitted to a single exponential equation to determine the corresponding relaxation time constants (**Supplementary Figs. 7, 8** and **Table 6**). The analysis was conducted using OriginPro 9.

### Proton-detection solid-state NMR experiments

2D hCH, 2D hChH, and 3D hCCH TOCSY correlation experiments were conducted using a 600 MHz Bruker Avance Neo spectrometer at Michigan State University, equipped with a 1.3 mm triple-resonance MAS probe. Samples were spun at 60 kHz MAS. The 2D hChH experiment employed a 0.8 ms RFDR-XY16 (radiofrequency-driven recoupling) mixing^72,73^. A total of 192 time-domain (TD) points were acquired for the indirect dimension, with 512 transients co-added per TD and a recycle delay of 2.0 s. The total experimental duration was 58 hr. A 2D hCH spectrum was acquired on the same 600 MHz Bruker spectrometer under identical conditions, but without RFDR mixing, to enable direct comparison. The hCH experiment were measured using a short second-CP of 100 µs. The 2D data were collected using the States-TPPI method^74^. Through-bond ^13^C-^13^C correlations were established using the 3D hCCH TOCSY (total correlation spectroscopy) experiment^75^, employing a 15 ms WALTZ-16 (wideband alternating-phase low-power technique for zero-residual splitting) mixing period at an rf field strength of 21.4 kHz^76^. A total of 128 × 128 TD points were acquired in the T_1_ and T_2_ evolution periods, with 8 transients co-added per TD point. The total experimental time was 78 h.

The Heteronuclear dipolar decoupling sequence slpTPPM (swept low-power two-pulse phase modulation)^77^ was applied during the T_1_ evolution period for the 2D hCH and 2D hChH sequences, and during both T_1_ and T_2_ periods in the 3D hCCH TOCSY experiment, with a rf field strength of 21.0 kHz. For the direct ^1^H detection period, WALTZ-16 decoupling was applied on the ^13^C channel with a rf field strength of 12.5 kHz for 2D hCH and 2D hChH, and at 21.4 kHz for 3D hCCH TOCSY. Water suppression was achieved using the MISSISSIPPI (multiple intense solvent suppression intended for sensitive spectroscopic investigation of protonated proteins) sequence^78^ on the ^1^H channel, with a rf of 15.2 kHz applied for 100 ms. The actual sample temperature was measured to be 304 K, based on the ^1^H chemical shift of water relative to the DSS signal at 0 ppm. ^1^H and ^13^C chemical shifts of *C. neoformans* polysaccharides, as well as ^13^C chemical shifts of melanin fragments detected in ^1^H-detected experiments, were documented in **Supplementary Tables 7** and **8**. Experimental details are summarized in **Supplementary Table 11**.

### MAS-DNP analysis of inter-polysaccharide interactions

A stock solution containing 10 mM AsymPol-POK bi-radicals in a d_6_-DMSO/H_2_O (10/90 vol%) mixture was prepared^48. 13^C, ^15^N-labeled *C. neoformans* H99 cells were mixed with 50 µL of the stock solution and gently ground using a set of mortar and pestle to ensure radical penetration and distribution into the porous cell walls. Approximately 30 mg of the processed sample was packed into a 3.2-mm sapphire rotor for DNP experiments. All experiments were conducted on a 600 MHz/395 GHz MAS-DNP spectrometer at the National High Magnetic Field Laboratory (Tallahassee, FL, USA), equipped with an 89 mm bore and a gyrotron microwave source. Data acquisition utilized a 3.2-mm HCN probe operating at 8 kHz MAS and 100 K. The gyrotron cathode current was maintained between 130-150 mA, with a voltage setting of 16.2 kV. The power of the microwave irradiation was 6.5 W. The NMR sensitivity enhancement (ε_on/off_) was 11, and the DNP signal buildup time was ∼2.9 s for *C. neoformans* carbohydrate signals. The PAR spectrum was acquired with 15 ms recoupling duration, during which the ^1^H and ^13^C irradiation frequencies were set at 53 kHz and 50 kHz, respectively^49,50^. The 2D ^13^C-^13^C DARR spectrum was recorded with a 100 ms mixing time.

## DATA AVAILABILITY

All relevant data that support the findings of this study are provided in the article and supplementary Information. All the original ssNMR data files will be deposited in the Zenodo repository, and the access code and DOI will be provided.

## AUTHOR CONTRIBUTIONS

A.A., I.G., and P.W. prepared labeled and unlabeled samples. A.A., I.G., D.R., and L.X. conducted ^13^C solid-state NMR experiments. J.R.Y. performed ^1^H-detection solid-state NMR experiments. F.J.S. and F. M.-V. performed DNP measurements. A.A., I.G., J.R.Y., C.C., R.E.S. and T.L.D. analyzed the data. P.W. and T.W. designed the experiments. All authors contributed to manuscript writing.

## COMPETING INTERESTS

The authors declare no competing interests.

## ACKNOWLEDGMENT

This work was primarily supported by the National Institutes of Health (NIH) grant R01AI173270 to T.W. A portion of this work was performed at the National High Magnetic Field Laboratory, which is supported by the National Science Foundation Cooperative Agreement No. DMR-2128556 and the State of Florida. The MAS-DNP system at NHMFL is funded in part by NIH RM1-GM148766. C.C. and R.E.S. acknowledge the support from NIH grant R01AI171093. F.S acknowledges a postdoctoral scholar award from the Provost’s Office at Florida State University.

## Notes

### Competing Interest Statement

The authors have declared no competing interest.

## REFERENCES

1 Mitchell, T. G. & Perfect, J. R. Cryptococcosis in the era of AIDS - 100 years after the discovery o Cryptococcus neoformans. Clinical Microbiol. Rev. 8, 515–548 (1995).

2 Kozel, T. R. Virulence factors of Cryptococcus neoformans. Trends Microbiol 3, 295–299 (1995).

3 Buchanan, K. L. & Murphy, J. W. What makes Cryptococcus neoformans a pathogen? Emerg. Infect. Dis. 4, 71–83 (1998).

4 Denning, D. W. Global incidence and mortality of severe fungal disease. Lancet Infect. Dis. 24, e428–438 (2024).

5 Brown, G. D. et al. Hidden Killers: Human Fungal Infections. Sci. Trans. Med. 4, 165rv113 (2012).

6 Cano, E., Yetmar, Z. & Razonable, R. Cryptococcus Species Other Than Cryptococcus neoformans and Cryptococcus gattii: Are They Clinically Significant? Open Forum Infect. Dis. 7, ofaa527 (2020).

7 Jackson, K., Ding, M. & Nielsen, K. Importance of Clinical Isolates in Cryptococcus neoformans Research. J. Fungi 9, 364 (2023).

8 Ballou, E. & Johnston, S. The cause and effect of Cryptococcus interactions with the host. Curr. Opin. Microbiol. 40, 88–94 (2017).

9 Tugume, L. et al. Cryptococcal meningitis. Nat. Rev. Dis. Primers 9, 62 (2023).

10 Rajasingham, R. et al. The global burden of HIV-associated cryptococcal infection in adults in 2020: a modelling analysis. Lancet Infect. Dis. 22, 1748–1755 (2022).

11 Meya, D. B. & Williamson, P. R. Cryptococcal Disease in Diverse Hosts. N. Engl. J. Med. 390, 1597–1610 (2024).

12 Chang, C. C. et al. Global guideline for the diagnosis and management of cryptococcosis: an initiative of the ECMM and ISHAM in cooperation with the ASM. Lancet Infect. Dis. 24, e495–512 (2024).

13 Denning, D. Echinocandins: a new class of antifungal. J. Antimicrob. Chemother. 49, 889–891 (2002).

14 Garcia-Rubio, R. et al. Critical Assessment of Cell Wall Integrity Factors Contributing to in vivo Echinocandin Tolerance and Resistance in Candida glabrata. Front. Microbiol. 12, 702779 (2021).

15 Mukaremera, L. The Cryptococcus wall: A different wall for a unique lifestyle. PLoS Pathog. 19, e1011141 (2023).

16 Garcia-Rubio, R., de Oliveira, H. C., Rivera, J. & Trevijano-Contador, N. The Fungal Cell Wall: Candida, Cryptococcus, and Aspergillus Species. Front. Microbiol. 10, 2993 (2019).

17 Doering, T. L. in Annu. Rev. Microbiol. Vol. 63 223–247 (2009).

18 Gilbert, N., Lodge, J. & Specht, C. Cryptococcus: From Human Pathogen to Model Yeast. (ASM Press, 2011).

19 Manners, D. J., Masson, A. J., Patterson, J. C., Bjorndal, H. & Lindberg, B. The structure of a beta-(1-6)-D-glucan from yeast cell walls. Biochem. J. 135, 31–36 (1973).

20 McClelland, E., Bernhardt, P. & Casadevall, A. Estimating the relative contributions of virulence factors for pathogenic microbes. Infect. Immun. 74, 1500–1504 (2006).

21 Agustinho, D., Miller, L., Li, L. & Doering, T. Peeling the onion: the outer layers of Cryptococcus neoformans. Mem. Inst. Oswaldo Cruz 113 (2019).

22 Fromtling, R., Shadomy, H. & Jacobson, E. Decreased virulence in stable, acapsular mutants of Cryptococcus neoformans. Mycopathologia 79, 23–29 (1982).

23 Casadevall, A., Rosas, A. & Nosanchuk, J. Melanin and virulence in Cryptococcus neoformans. Curr. Opin. Microbiol. 3, 354–358 (2000).

24 Casadevall, A. et al. The capsule of Cryptococcus neoformans. Virulence 10, 822–831 (2019).

25 Bose, I., Reese, A., Ory, J., Janbon, G. & Doering, T. A yeast under cover: the capsule of Cryptococcus neoformans. Eukaryot. Cell 2, 655–663 (2003).

26 Cherniak, R. & Sundstrom, J. Polysaccharide Antigens of the Capsule of Cryptococcus neoformans. Infect. Immun. 62, 1507–1512 (1994).

27 Frases, S., Nimrichter, L., Viana, N., Nakouzi, A. & Casadevall, A. Cryptococcus neoformans capsular polysaccharide and exopolysaccharide fractions manifest physical, chemical, and antigenic differences. Eukaryot. Cell 7, 319–327 (2008).

28 Camacho, E. et al. The structural unit of melanin in the cell wall of the fungal pathogen Cryptococcus neoformans. J. Biol. Chem. 294, 10471–10489 (2019).

29 Baker, R., Chrissian, C., Stark, R. & Casadevall, A. Cryptococcus neoformans melanization incorporates multiple catecholamines to produce polytypic melanin. J. Biol. Chem. 298, 101519 (2022).

30 Ikeda, R., Sugita, T., Jacobson, E. & Shinoda, T. Effects of melanin upon susceptibility of Cryptococcus to antifungals. Microbiol. Immunol. 47, 271–277 (2003).

31 Wang, Y., Aisen, P. & Casadevall, A. Cryptococcus neoformans melanin and virulence: mechanism of action. Infect. Immun. 63, 3131–3136 (1995).

32 Wang, Y., Aisen, P. & Casadevall, A. Melanin, melanin ‘‘ghosts,’’ and melanin composition in Cryptococcus neoformans. Infect. Immun. 64, 2420–2424 (1996).

33 Ni, Q. Z. et al. High frequency dynamic nuclear polarization. Acc. Chem. Res. 46, 1933–1941 (2013).

34 Chow, W. Y., De Paëpe, G. & Hedinger, S. Biomolecular and Biological Applications of Solid-State NMR with Dynamic Nuclear Polarization Enhancement. Chem. Rev. 122, 9795–9847 (2022).

35 Rossini, A. J. et al. Dynamic nuclear polarization surface enhanced NMR spectroscopy. Acc. Chem. Res. 46, 1942–1951 (2013).

36 Lee, D., Hediger, S. & De Paege, G. Is solid-state NMR enhanced by dynamic nuclear polarization? Solid State Nucl. Magn. Reson. 66, 6–20 (2015).

37 Evans, E. The Antigenic Composition of Cryptococcus Neoformans: I. A Serologic Classification by Means of the Capsular and Agglutination Reactions. J. Immunol. 64, 423–430 (1950).

38 Wilson, D., Bennett, J. & Bailey, J. Serologic grouping of Cryptococcus neoformans. Proc. Soc. Exp. Biol. Med. 127, 820–823 (1968).

39 Franzot, S., Salkin, I. & Casadevall, A. Cryptococcus neoformans var. grubii: Separate varietal status for Cryptococcus neoformans serotype A isolates. J. Clin. Microbiol. 37, 838–840 (1999).

40 Kwon-Chung, K. & Varma, A. Do major species concepts support one, two or more species within Cryptococcus neoformans? FEMS Yeast Res. 6, 574–587 (2006).

41 Hu, G. et al. Transcriptional Regulation by Protein Kinase A in Cryptococcus neoformans. PLoS Pathog. 3, e42 (2007).

42 Fromtling, R. A., Shadomy, H. J. & Jacobson, E. S. Decreased virulence in stable, acapsular mutants of Cryptococcus neoformans. Mycopathologia 79, 23–29 (1982).

43 Hou, G., Yan, S., Trébosc, J., Amoureux, J. & Polenova, T. Broadband homonuclear correlation spectroscopy driven by combined R2nv sequences under fast magic angle spinning for NMR structural analysis of organic and biological solids. J. Magn. Reson. 232, 18–30 (2013).

44 Lends, A. et al. Molecular Distinction of Cell Wall and Capsular Polysaccharides in Encapsulated Pathogens by In Situ Magic-Angle Spinning NMR Techniques. J. Am. Chem. Soc. 147, 6813–6824 (2025).

45 Ader, C. et al. Structural rearrangements of membrane proteins probed by water-edited solid-state NMR spectroscopy. J. Am. Chem. Soc. 131, 170–176 (2009).

46 Luo, W. & Hong, M. Conformational Changes of an Ion Channel Detected Through Water-Protein Interactions Using Solid-State NMR Spectroscopy. J. Am. Chem. Soc. 132, 2378–2384 (2010).

47 White, P. B., Wang, T., Park, Y. B., Cosgrove, D. J. & Hong, M. Water–polysaccharide interactions in the primary cell wall of Arabidopsis thaliana from polarization transfer solid-state NMR. J. Am. Chem. Soc. 136, 10399–10409 (2014).

48 Mentink-Vigier, F. et al. Computationally assisted design of polarizing agents for dynamic nuclear polarization enhanced NMR: The asymPol family. J. Am. Chem. Soc. 140, 11013–11019 (2018).

49 De Paëpe, G., Lewandowski, J. R., Loquet, A., Bockmann, A. & Griffin, R. G. Proton assisted recoupling and protein structure determination. J. Chem. Phys. 129, 245101 (2008).

50 Donovan, K. J., Jain, S. K., Silvers, R., Linse, S. & Griffin, R. G. Proton-Assisted Recoupling (PAR) in Peptides and Proteins. J. Phys. Chem. B 121, 10804–10817 (2017).

51 Chatterjee, S. et al. Using Solid-State NMR To Monitor the Molecular Consequences of Cryptococcus neoformans Melanization with Different Catecholamine Precursors. Biochem. 51, 6080–6088 (2012).

52 Cao, W. et al. Unraveling the Structure and Function of Melanin through Synthesis. J. Am. Chem. Soc. 143, 2622–2637 (2021).

53 Chatterjee, S. et al. The melanization road more traveled by: Precursor substrate effects on melanin synthesis in cell-free and fungal cell systems. J. Biol. Chem. 293, 20157–20168 (2018).

54 Chrissian, C. et al. Melanin deposition in two Cryptococcus species depends on cell-wall composition and flexibility. J. Biol. Chem. 295, 1815–1828 (2020).

55 Eisenman, H. C., Chow, S.-K., Tse, K. K., McClelland, E. E. & Casadevall, A. The effect of L-DOPA on Cryptococcus neoformans growth and gene expression. Virulence 2, 329–336 (2011).

56 Safeer, A. et al. Probing Cell-Surface Interactions in Fungal Cell Walls by High-Resolution 1H-Detected Solid-State NMR Spectroscopy. Chem. Eur. J. 29, e202202616 (2023).

57 Kang, X. et al. Molecular architecture of fungal cell walls revealed by solid-state NMR. Nat. Commun. 9, 2747 (2018).

58 Lamon, G. et al. Solid-state NMR molecular snapshots of Aspergillus fumigatus cell wall architecture during a conidial morphotype transition. Proc. Natl. Acad. Sci. USA 120, e2212003120 (2023).

59 Fernando, L. D. et al. Structural adaptation of fungal cell wall in hypersaline environment. Nat. Commun. 14, 7082 (2023).

60 Ehren, H. L. et al. Characterization of the cell wall of a mushroom forming fungus at atomic resolution using solid-state NMR spectroscopy. Cell Surf. 6, 100046 (2020).

61 Ghassemi, N. et al. Solid-State NMR Investigations of Extracellular Matrixes and Cell Walls of Algae, Bacteria, Fungi, and Plants. Chem. Rev. 122, 10036–10086 (2022).

62 Latgé, J. P. & Wang, T. Modern Biophysics Redefines Our Understanding of Fungal Cell Wall Structure, Complexity, and Dynamics. mBio 13, e0114522 (2022).

63 Mukaremera, L. et al. Titan cell production in Cryptococcus neoformans reshapes the cell wall and capsule composition during infection. Cell Surf. 1, 15–24 (2018).

64 Bekirian, C. et al. β-1,6-glucan plays a central role in the structure and remodeling of the bilaminate fungal cell wall. eLife 13, RP100569 (2024).

65 Reese, A. J. & Doering, T. L. Cell wall alpha-1,3-glucan is required to anchor the Cryptococcus neoformans capsule. Mol. Microbiol. 50, 1401–1409 (2003).

66 Lesage, A., Bardet, M. & Emsley, L. Through-bond carbon-carbon connectivities in disordered solids by NMR. J. Am. Chem. Soc. 121, 10987–10993 (1999).

67 Cheng, Q. et al. Molecular architecture of chitin and chitosan-dominated cell walls in zygomycetous fungal pathogens by solid-state NMR. Nat. Commun. 15, 8295 (2024).

68 Dickwella Widanage, M. C. et al. Adaptative survival of Aspergillus fumigatus to echinocandins arises from cell wall remodeling beyond β-1,3-glucan synthesis inhibition. Nat. Commun. 15, 6382 (2024).

69 Torchia, D. Measurement of proton-enhanced C-13 T1 values by a method which suppresses artifacts. J. Magn. Reson. 30, 613–616 (1978).

70 van Rossum, B., de Groot, C., Ladizhansky, V., Vega, S. & de Groot, H. A method for measuring heteronuclear (1H-13C) distances in high speed MAS NMR. J. Am. Chem. Soc. 122, 3465–3472 (2000).

71 Huster, D., Xiao, L. & Hong, M. Solid-state NMR investigation of the dynamics of the soluble and membrane-bound colicin Ia channel-forming domain. Biochemistry 40, 7662–7674 (2001).

72 Bennett, A. E., Griffin, R. G., Ok, J. H. & Vega, S. Chemical shift correlation spectroscopy in rotating solids: Radio frequency-driven dipolar recoupling and longitudinal exchange. J. Chem. Phys. 96, 8624–8627 (1992).

73 Nishiyama, Y., Zhang, R. & Ramamoorthy, A. Finite-pulse radio frequency driven recoupling with phase cycling for 2D 1H/1H correlation at ultrafast MAS frequencies. J. Magn. Reson. 243, 25–32 (2014).

74 Marion, D., Ikura, M., Tschudin, R. & Bax, A. D. Rapid recording of 2D NMR spectra without phase cycling. Application to the study of hydrogen exchange in proteins. J. Magn. Reson. 85, 393–399 (1989).

75 Andreas, L. B. et al. Structure of fully protonated proteins by proton-detected magic-angle spinning NMR. Proc. Natl. Acad. Sci. USA 113, 9187–9192 (2016).

76 Shaka, A., Keeler, J., Frenkiel, T. & Freeman, R. An improved sequence for broadband decoupling: WALTZ-16. J. Magn. Reson. 52, 335–338 (1983).

77 Lewandowski, J. R. et al. Measurement of Site-Specific 13C Spin-Lattice Relaxation in a Crystalline Protein. J. Am. Chem. Soc. 132, 8252–8254 (2010).

78 Zhou, D. H. & Rienstra, C. M. High-performance solvent suppression for proton detected solid-state NMR. J. Magn. Reson. 192, 167–172 (2008).

