## Supplementary Material for "Polymorphic α-Glucans as Structural Scaffolds in *Cryptococcus* Cell Walls for Chitin, Capsule, and Melanin: Insights from ^13^C and ^1^H Solid-State NMR"

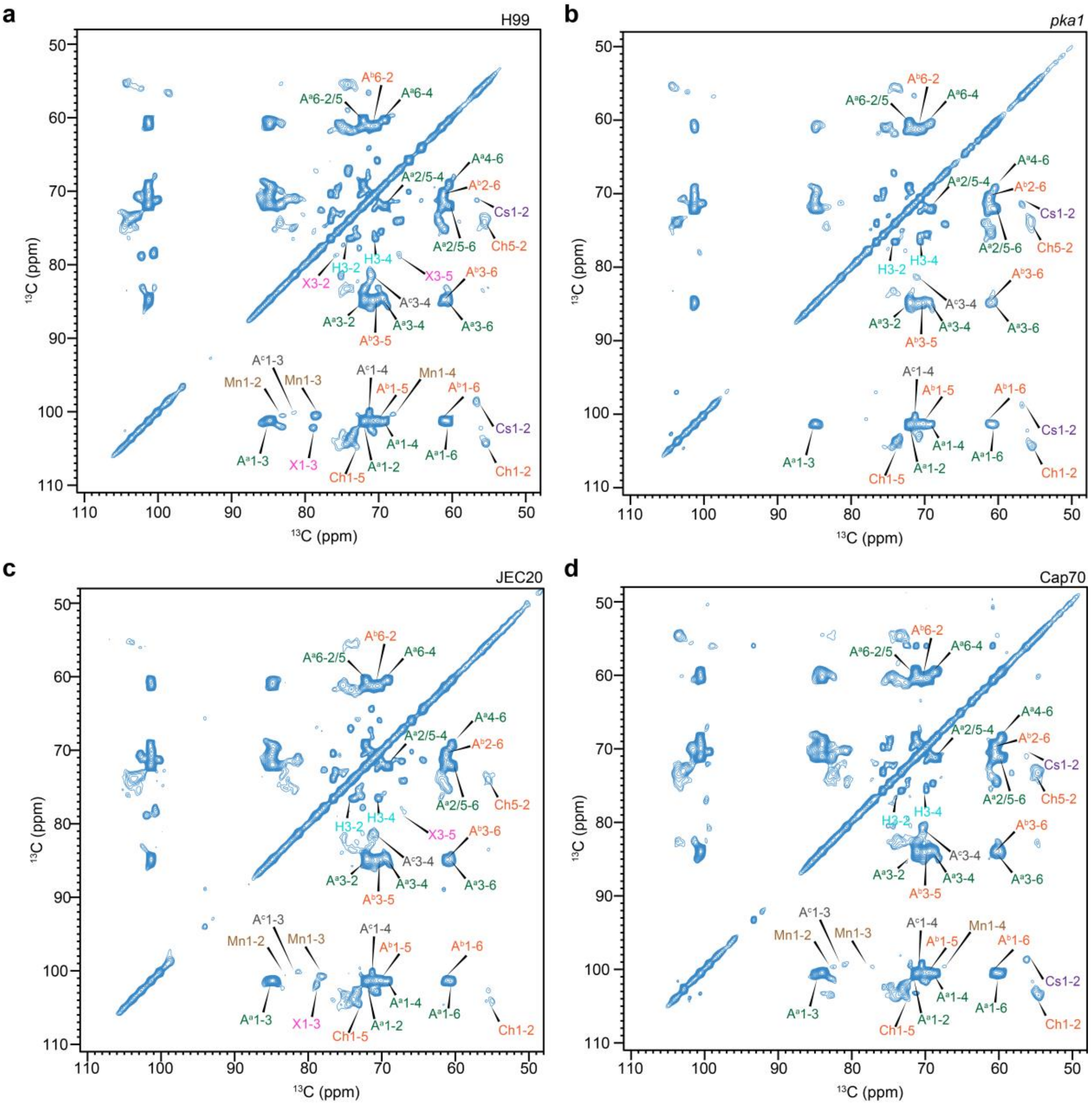

**Supplementary Figure 1. Resonance assignment of rigid glucans in *C. neoformans* cell wall.** CP-based 2D $^{13}C$-$^{13}C$ correlation spectrum measured with 53 ms CORD mixing for **(a)** H99, **(b)** *pka1*, **(c)** JEC20, and **(d)** Cap70.

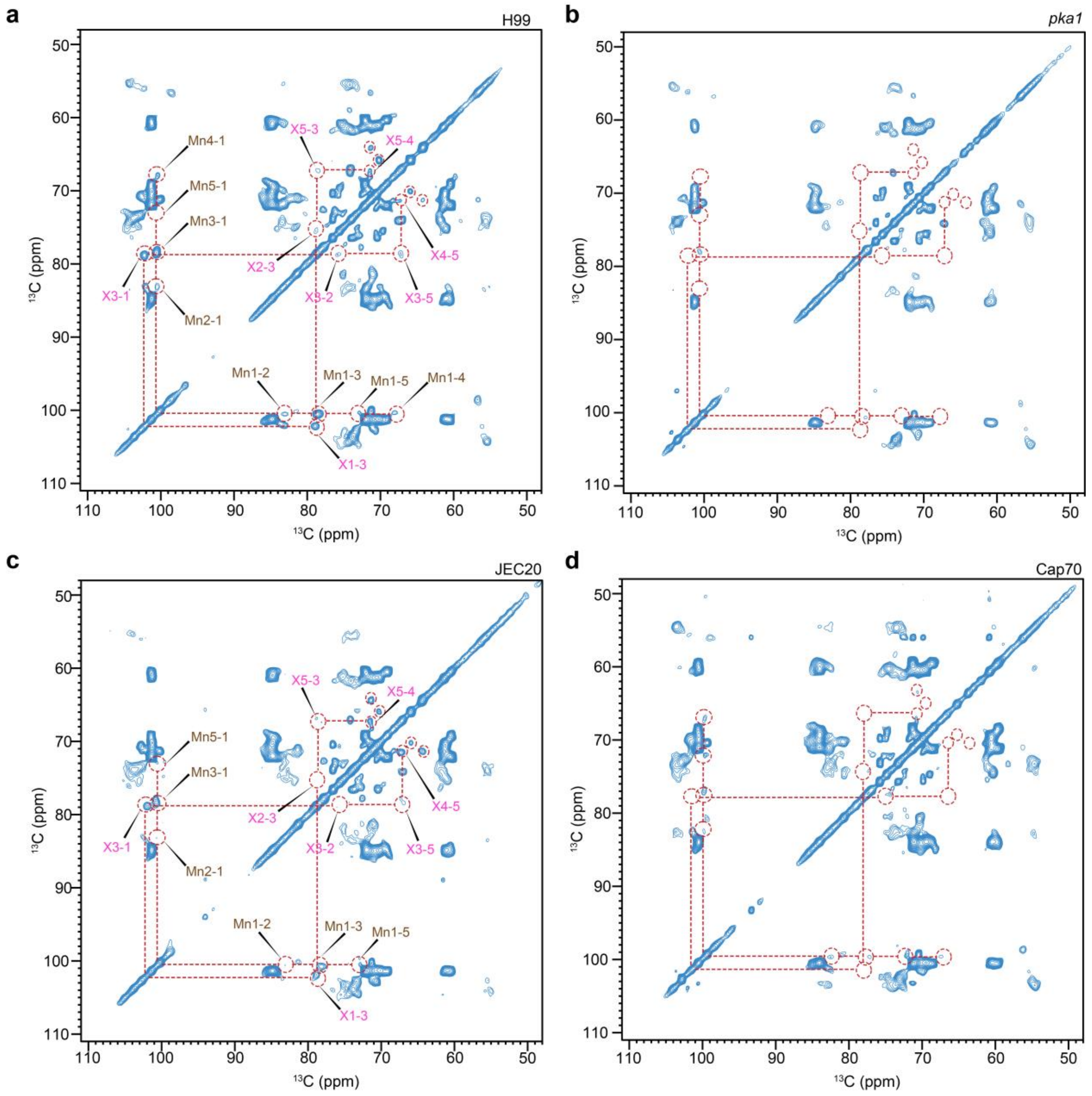

**Supplementary Figure 2. Resonance assignment of capsular molecules in *C. neoformans*. (a)** CP-based 2D $^{13}$C-$^{13}$C correlation spectrum measured with 53 ms CORD mixing for H99, highlighting mannan and xylose signals arising from capsules. **(b)** CP-based 2D $^{13}$C-$^{13}$C correlation spectrum measured with 53 ms CORD mixing for *pka1* highlighting the absence of mannan and xylose signals. **(c)** CP-based 2D $^{13}$C-$^{13}$C correlation spectrum measured with 53 ms CORD mixing for JEC20 highlighting mannan and xylose signals arising from capsules. **(d)** CP-based 2D $^{13}$C-$^{13}$C correlation spectrum measured with 53 ms CORD mixing for Cap70 highlighting the presence of mannan and the absence of xylose signals.

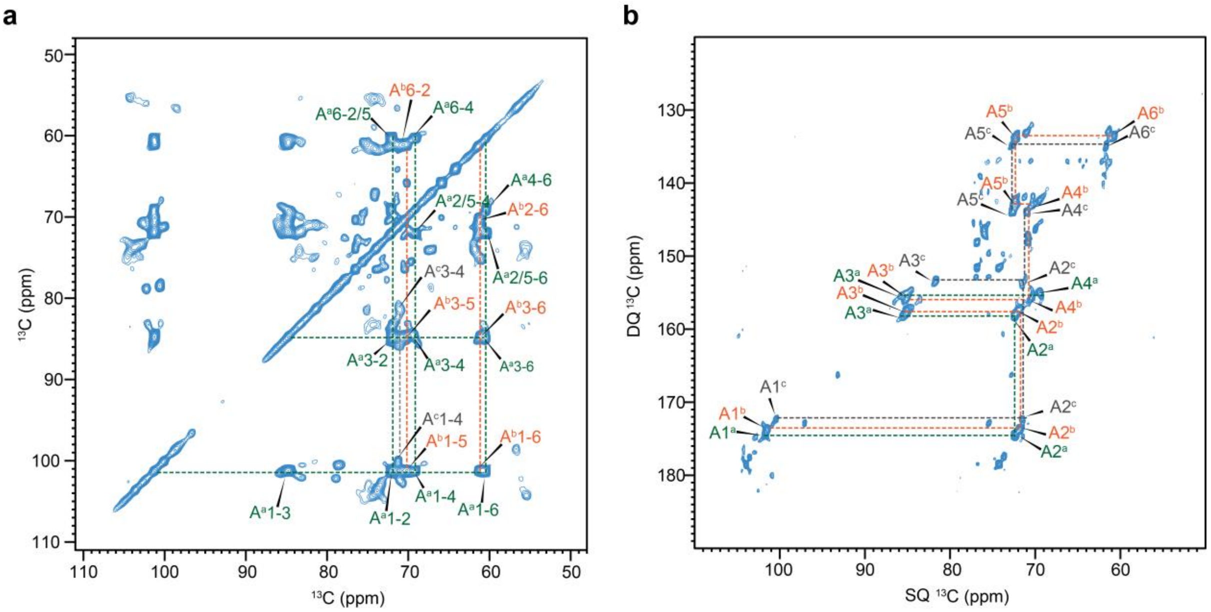

**Supplementary Figure 3. Resonance assignment of α-1,3-glucan in *C. neoformans* cell wall. (a)** CP-based 2D $^{13}$C-$^{13}$C correlation spectrum measured with 53 ms CORD mixing. **(b)** Through-bond carbon connectivity was resolved using 2D $^{13}$C CP refocused J-INADEQUATE spectrum. Multiple sets of conformers were resolved for α-1,3-glucan. Each peak is annotated with the abbreviation of the carbohydrate name, the subtype (in superscript), and the carbon number. For instance, A$^c$1 represents the carbon 1 of type-c α-1,3-glucan.

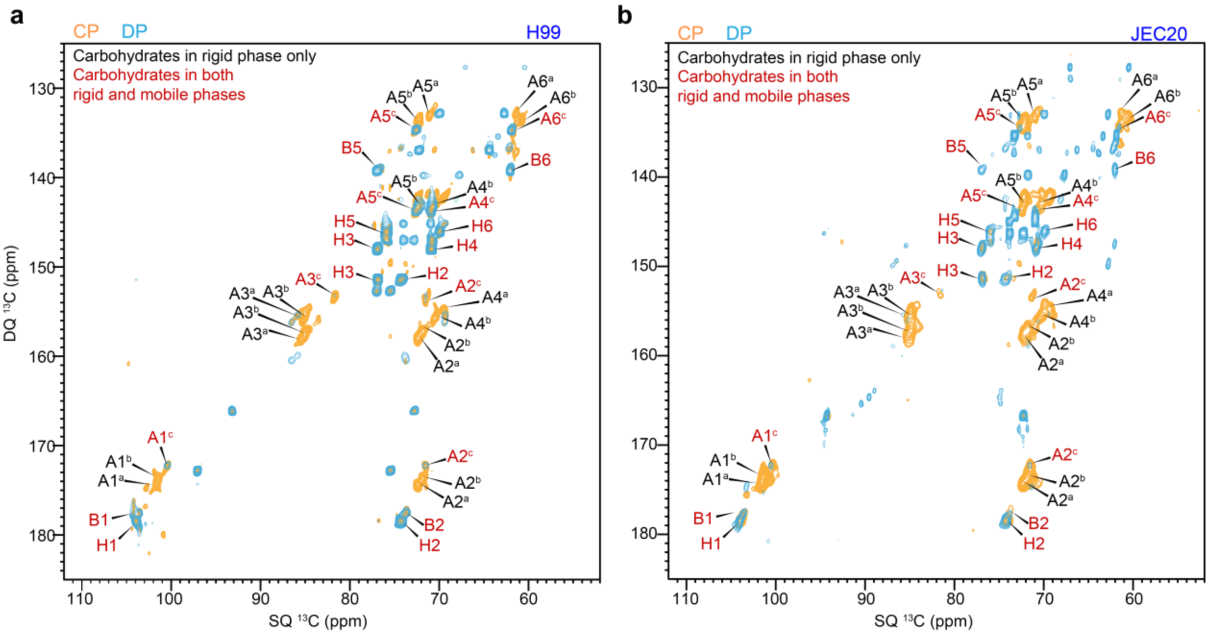

**Supplementary Figure 4. Distribution of α-1,3-glucan, and β-glucan in rigid and mobile domains.** Overlay of 2D refocused J-INADEQUATE spectra measured with CP (orange) and DP (cyan) for **(a)** *C. neoformans* H99 and **(b)** *C. neoformans* JEC20 samples. The carbohydrates observed only in the CP-based spectra are rigid and are marked in black. The carbohydrates observed in both CP (INADEQUATE here and CORD in Figure S1) and DP-based spectra are marked in red: these carbohydrates have two-modal distribution in rigid and mobile phases. β-1,6-glucan, β-1,3-glucan, and types-c α-1,3-glucan were observed in both domains for both samples.

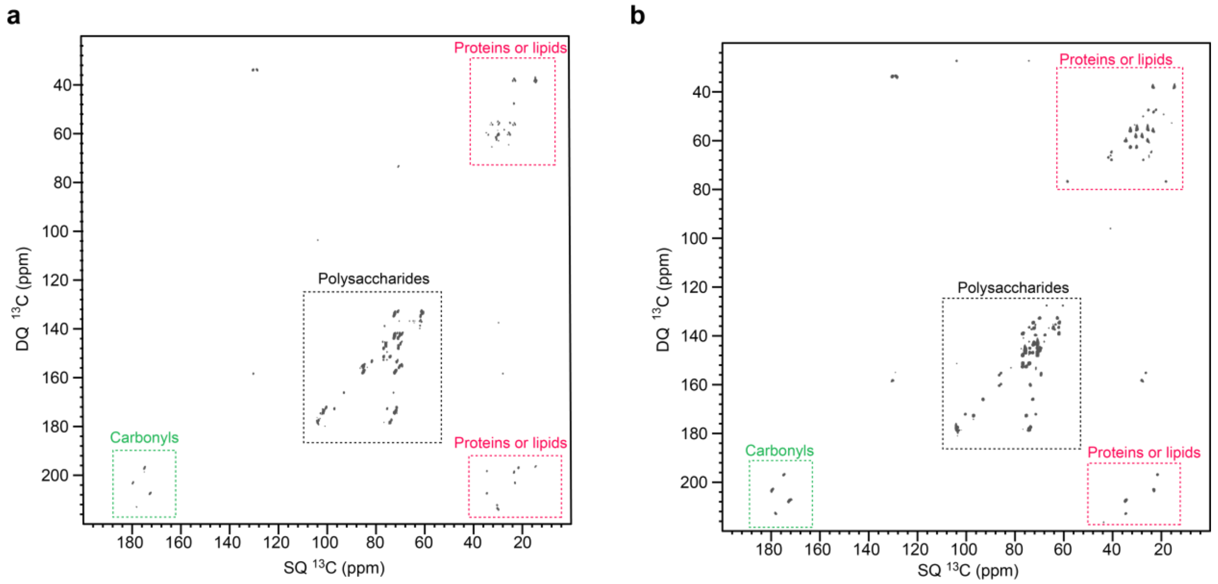

**Supplementary Figure 5. Biopolymer signals resolved in 2D $^{13}$C refocused J-INADEQUATE. (a)** Rigid components of the cell wall in *C. neoformans* sample probed using 2D CP $^{13}$C refocused J-INADEQUAT E spectrum. The individual regions were marked with quadrangles of different colors: green for carbonyls, black for polysaccharides, and red for aliphatics from proteins and lipids. **(b)** Mobile components of *C. neoformans* sample detected using 2D DP $^{13}$C refocused J-INADEQUATE spectrum. The individual regions are marked using the same color code mentioned above.

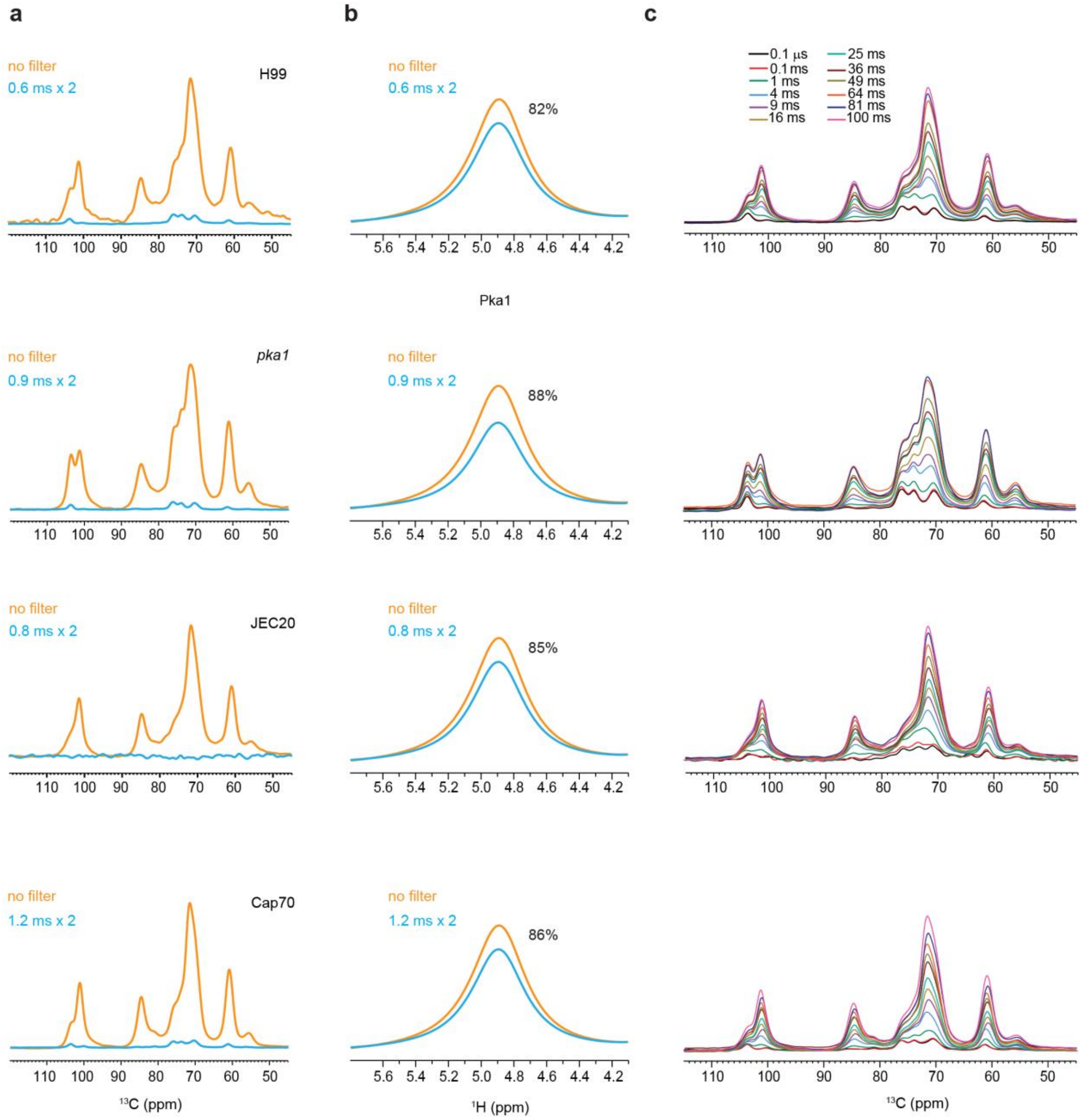

**Supplementary Figure 6. Water-edited experiment setup for inspecting carbohydrate hydration. (a)** $^{1}$H-T$_2$ filtered (blue) and control (orange) $^{13}$C spectra are shown for four strains. From the top to bottom: H99, *pka1*, JEC20, and Cap70. No spin diffusion was applied. Approximately 85% of carbohydrate $^{13}$C signals were removed by the T$_2$ filter. **(b)** $^{1}$H-T$_2$ filtered (blue) and control (orange) $^{1}$H NMR spectra, with 82% and 88% of water signal retained for each strain, after the $^{1}$H T$_2$ filter. From the top to bottom: H99, *pka1*, JEC20, and Cap70. **(c)** 1D water-edited $^{13}$C spectra with different $^{1}$H mixing times. From the top to bottom: H99, *pka1*, JEC20, and Cap70. All spectra were measured on a 400 MHz spectrometer at 15 kHz MAS at 280 K.

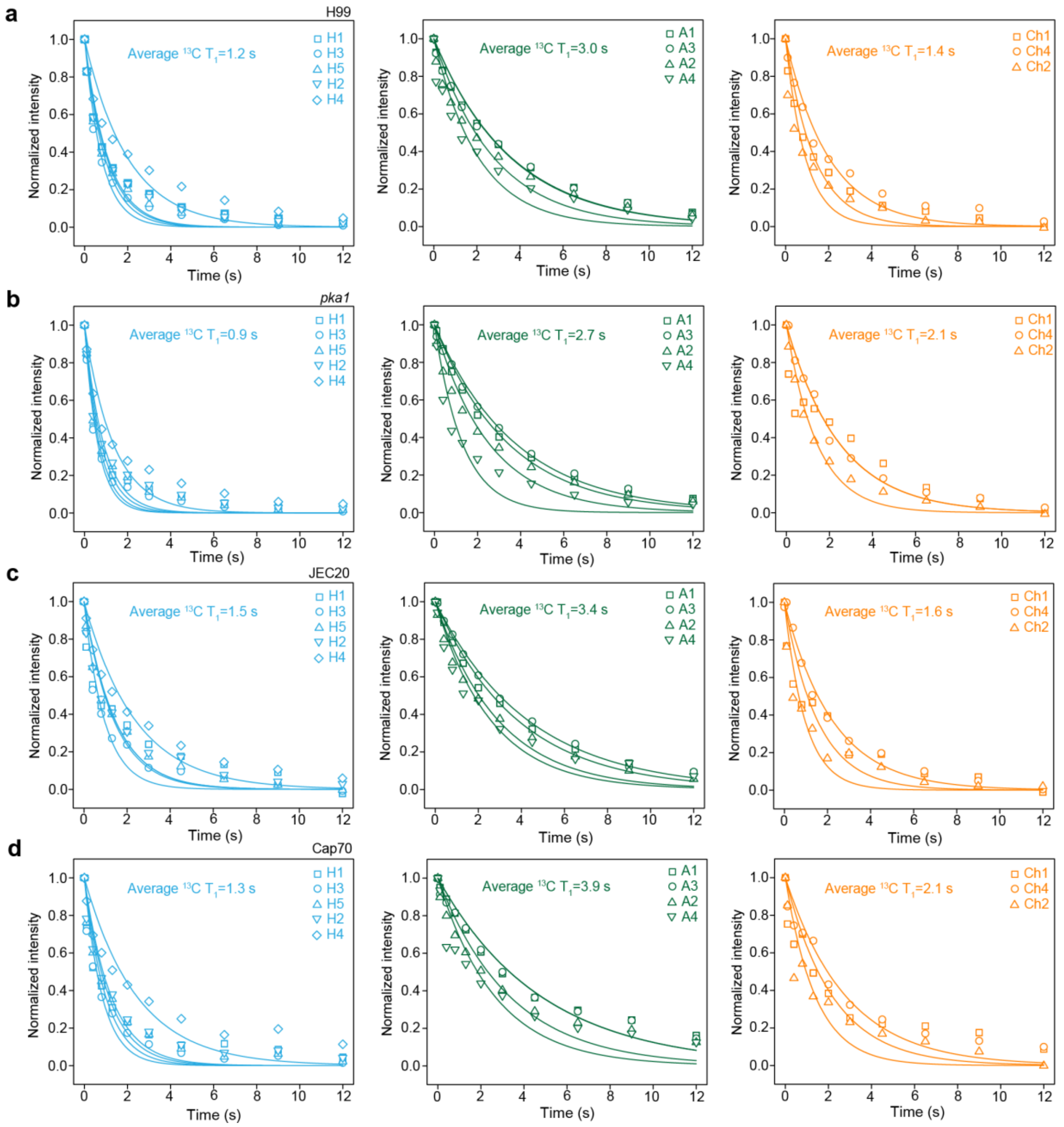

**Supplementary Figure 7. $^{13}$C-$T_1$ relaxation of polysaccharides in *C. neoformans*.** $^{13}$C-$T_1$ measured with Torchia CP for **(a)** H99 **(b)** *pka1* **(c)** JEC20 and **(d)** Cap70 samples. The data are separately presented for β-1,6-glucan (light blue), α-1,3-glucan (green), and chitin (orange). The acquired data were fitted to a single exponential decay equation. Different symbols and color codes are used to represent different carbons in these polysaccharides.

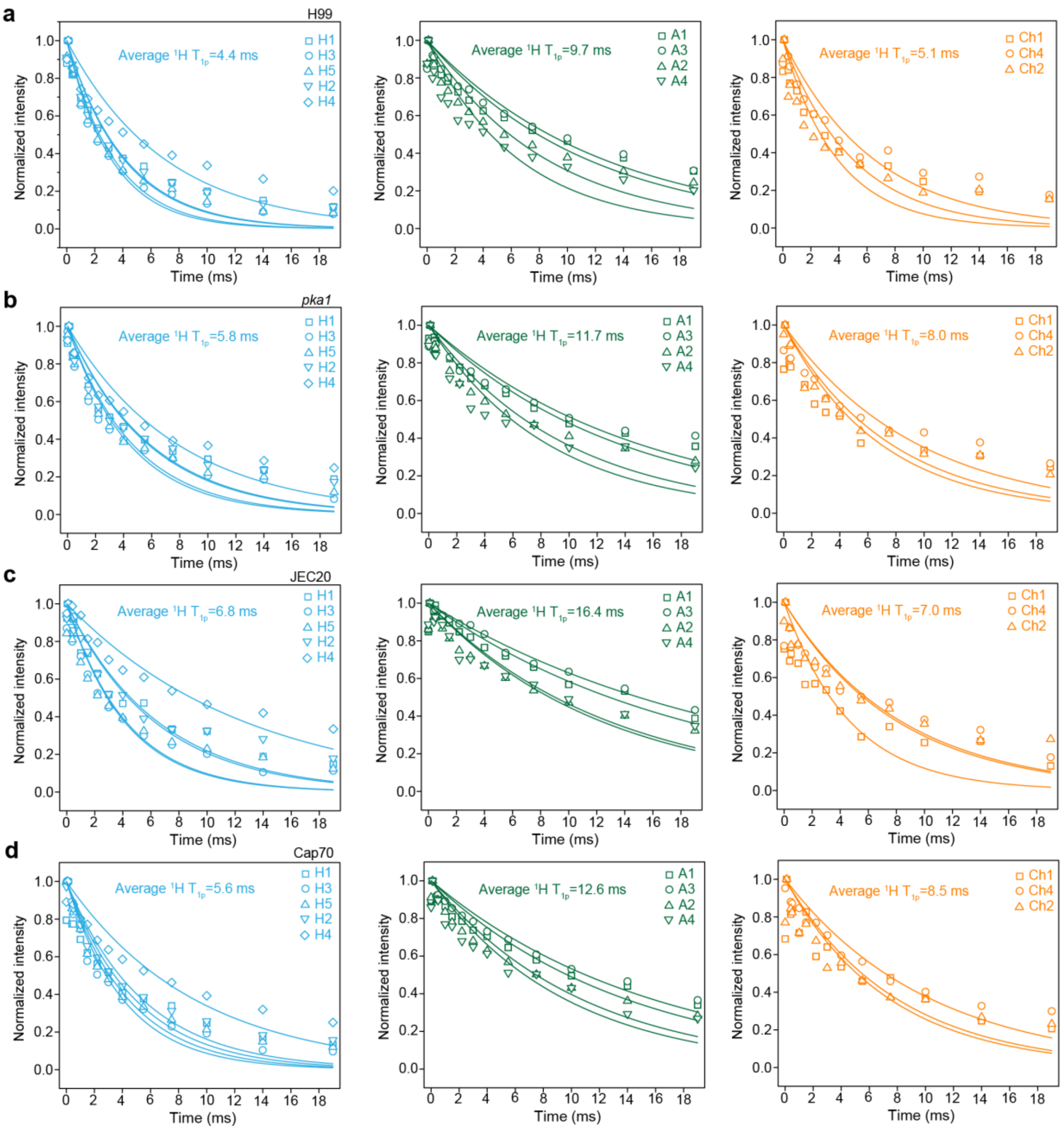

**Supplementary Figure 8. $^{1}$H-T$_{1\rho}$ relaxation of polysaccharides in *C. neoformans*.** $^{1}$H-T$_{1\rho}$ measured with Torchia CP for **(a)** H99 **(b)** *pka1* **(c)** JEC20 and **(d)** Cap70 samples. The data are separately presented for β-1,6-glucan (light blue), α-1,3-glucan (green), and chitin (orange). The acquired data were fitted to a single exponential decay equation. Different symbols and color codes are used to represent different carbons in these polysaccharides.

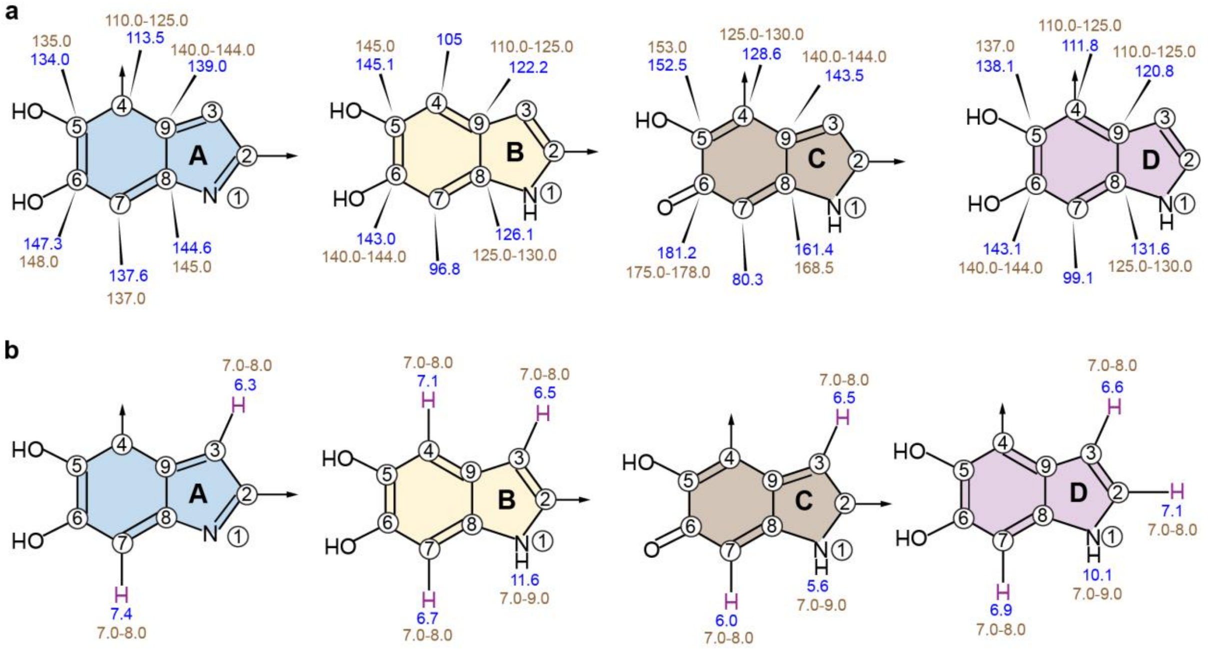

**Supplementary Figure 9. Chemical shifts of melanin fragments in *C. neoformans*. (a)** $^{13}$C and **(b)** $^{1}$H chemical shifts of melanin fragments. Chemical shifts labeled in brown are melanin signals experimentally observed in fast-MAS 2D hChH spectrum. Chemical shifts labeled in blue are predicted by ChemDraw 23.1.1 software.

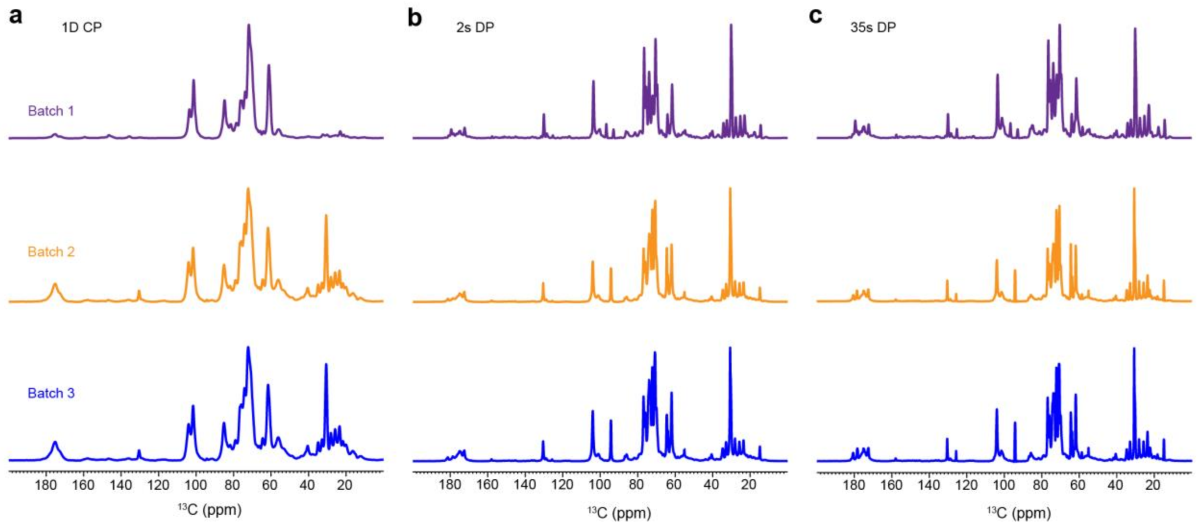

**Supplementary Figure 10. Replication of *C. neoformans* samples.** (**a**) 1D $^{13}$C CP spectra of *C. neoformans*. (**b**) Mobile components detected by 1D $^{13}$C DP with a short recycle delay of 2 s. (**c**) Quantitative $^{13}$C DP spectra for the detection of all molecules through a recycle delay of 30 s. The spectra exhibit high reproducibility across three separate batches prepared in 2023 (batch 1), 2024 (batch 2), and 2025 (batch 3). The spectra are highly reproducible except for signals of unused glucose at 93 ppm.

**Supplementary Table 1. The average cell thickness of *Cryptococcus* cells.** Results are the average and standard deviation of measurements from different cells (n=24 for H99 and *pka1*, n=19 for JEC20, and n=32 for Cap70).  for each sample. A source data file is provided to document each single reading. n=number of cells

| Sample | Strain | Average cell thickness (µm) |
|---|---|---|
| *C. neoformans* | H99 | 5.7±0.5 |
| | *pka1* | 4.6±0.7 |
| | JEC20 | 5.9±0.6 |
| | Cap70 | 4.2±0.3 |

**Supplementary Table 2. $^{13}$C chemical shifts of *C. neoformans* polysaccharides in cell walls from $^{13}$C-based experiments**. The chemical shifts are references on the TMS scale. All chemical shifts are from room-temperature experiments, except for those underlined, which are from DNP-enhanced experiments.

| Carbohydrates | form | C1 | C2 | C3 | C4 | C5 | C6 | Reference |
|---|---|---|---|---|---|---|---|---|
| Rigid molecules | | | | | | | | |
| α-1,3-glucan (A) | a | 101.6 | 72.4 | 85.5 | 69.4 | 71.2 | 61.4 | Chakraborty *et al.* 2021[1] |
| | | 101.2 | 70.6 | 85.5 | / | 70.6 | 61.2 | |
| | b | 101.4 | 71.8 | 85.0 | 70.5 | 72.2 | 60.7 | |
| | | 101.0 | 71.6 | 84.5 | / | / | 60.0 | |
| | c | 100.4 | 71.5 | 81.9 | 71.1 | 72.6 | 61.6 | |
| | d | 100.5 | 70.2 | 87.6 | 67.8 | 70.2 | 59.9 | |
| β-1,6-glucan (H) | | 103.8 | 74.2 | 76.8 | 70.8 | 75.8 | 69.8 | Lowman *et al.* 2011[2] |
| β-1,3-glucan (B) | a | 103.6 | 73.7 | 86.5 | 69.3 | 76.8 | 62.0 | Chakraborty *et al.* 2021[1]<br>Shim *et al.* 2007[3]<br>Fairweather *et al.* 2009[4]<br>Saito *et al.* 1979[5] |
| | b | 103.6 | 73.7 | 85.8 | 69.3 | 76.8 | 62.0 | |
| Chitin | | 104.3 | 55.3 | 73.7 | 83.0 | 75.6 | 62.0 | Kang *et al.* 2018[6]<br>Fernando *et al.* 2021[7] |
| | | 103.4 | 54.5 | 73.9 | 82.5 | 73.9 | 62.0 | |
| Chitosan | | 98.4 | 56.6 | / | / | / | / | |
| | | 99.1 | 55.5 | / | 80.2 | / | 60.7 | |
| α-1,2,3-Mannan (Mn[1,2,3]) | | 100.5 | 83.0 | 78.5 | 68.0 | 72.8 | n.d. | Bacon *et al.* 1995[8]<br>Moyrand *et al.* 2002[9] |
| Xylose | | 102.2 | 75.8 | 78.8 | 71.3 | 67.1 | n.a. | |
| Mobile molecules | | | | | | | | |
| β-1,6-glucan (H) | | 103.8 | 74.2 | 76.8 | 70.8 | 75.8 | 69.8 | Lowman *et al.* 2011[2] |
| β-1,3-glucan (B) | a | 103.6 | 73.7 | 86.5 | 69.3 | 76.8 | 62.0 | Chakraborty *et al.* 2021[1]<br>Shim *et al.* 2007<br>Fairweather *et al.* 2009[4]<br>Saito *et al.* 1979[5] |
| | b | 103.6 | 73.7 | 85.8 | 69.3 | 76.8 | 62.0 | |
| α-1,3-glucan (A) | a | / | / | / | / | / | / | |
| | b | / | / | / | / | / | / | |
| | c | 100.4 | 71.5 | 82.0 | 71.1 | 72.6 | 61.6 | |
| α-1,2-Mannan (Mn[1,2]) | | 101.4 | 79.2 | 71.7 | 67.7 | 74.3 | 62.1 | Chakraborty *et al.* 2021[1] |

**Supplementary Table 3. The molar composition of rigid polysaccharides.** The numbers are estimated using integrals (volume) of cross peaks in 2D $^{13}C$-$^{13}C$ 53 ms CORD spectra. The average integrals of cross-peaks of each polysaccharide are shown. Error bars are standard errors. Chitosan signals in JEC20 cells have low intensity.

| Polysaccharide | | | | | | | | |
|---|---|---|---|---|---|---|---|---|
| Strain | α-1,3-glucan | | | β-1,6-glucan | Chitin | Chitosan | Mannan | Xylose |
| | a | b | c | | | | | |
| *C. neoformans* H99 | 41±13 | 20±5 | 9±5 | 8±2 | 4±0.7 | 2±0.6 | 11±5 | 5±2 |
| *C. neoformans pka1* | 34±9 | 21±3 | 11±5 | 21±5 | 8±1.0 | 5±0.6 | / | / |
| *C. neoformans* JEC20 | 38±11 | 28±8 | 6±2 | 7±3 | 5±2 | / | 6±3 | 10±4 |
| *C. neoformans* Cap70 | 39±15 | 36±6 | 9±4 | 7±2 | 5±0.7 | 2±0.6 | 2±0.4 | / |

The area of the following well-resolved cross peaks 53 ms CORD spectra are used:
α-1,3 (a): the average of C1-C2/3/4 and C3-C2/4.
α-1,3 (b): the average of C1-C2/4, C3-2/4.
α-1,3 (c): the average of C1-C2/4, C3-2/4.
β-1,6: the average of C3-C2/4, C5-C4/6.
Chitin: the average of C1-2/5, C5-C2.
Chitosan: the average of C1-2, C3-C2.
Mannan[1,2,3]: the average of C1-C2/3/4/5.
Xylose: the average of C1-C3, C3-2/5, C4-5.

**Supplementary Table 4. The molar composition of mobile polysaccharides.** The numbers are estimated using integrals (volume) of cross peaks in 2D $^{13}C$-$^{13}C$ refocused DP-J INADEQUATE spectra. The average integrals of cross-peaks of each polysaccharide are shown. Error bars are standard errors of the peak integrals.

| Polysaccharide | | | | |
|---|---|---|---|---|
| Strains | β-1,6-glucan | β-1,3-glucan | α-1,3-glucan (c) | Mannan |
| *C. neoformans* H99 | 71±11 | 21±9 | 4±1.0 | 4±0.9 |
| *C. neoformans pka1* | 77±13 | 17±5 | 3±0.5 | 3±0.7 |
| *C. neoformans* JEC20 | 56±11 | 15±6 | 7±2 | 22±6 |
| *C. neoformans* Cap70 | 68±13 | 19±8 | 4±1.0 | 9±1.6 |

The area of the following well-resolved cross peaks refocused DP-J INADEQUATE spectra are used:
β-1,6: the average of C1, C2, C3, C4, C5, and C6.
β-1,3; the average of C1, C2, C3, C4, C5, and C6.
α-1,3-glucan: the average of C1, C2, and C3.
Mannan[1,2]: the average of C3, C4, C5 and C6.

**Supplementary Table 5. Water-edited intensities of polysaccharides.** Intensity ratios are obtained by comparing the peak intensities in water-edited and control spectra. The average values for each molecule in each sample are highlighted. Error bars are s.d. propagated from NMR signal-to-noise ratios.

| Polysaccharide | Cross-peak | *C. neoformans* (Serotype A) | | *C. neoformans* (Serotype D) | |
|---|---|---|---|---|---|
| | | H99 | *pka1* | JEC20 | Cap70 |
| α-1,3-glucan ($A^a$) | A1-1 | 0.11±0.06 | 0.31±0.03 | 0.33±0.04 | 0.37±0.02 |
| | A1-3 | 0.58±0.08 | 0.15±0.08 | 0.39±0.01 | 0.31±0.06 |
| | A1-2/5 | 0.29±0.02 | 0.24±0.04 | 0.30±0.05 | 0.42±0.03 |
| | A1-A4 | 0.31±0.03 | 0.28±0.06 | 0.02±0.01 | 0.48±0.05 |
| | A3-A1 | 0.34±0.08 | 0.19±0.05 | 0.33±0.02 | 0.32±0.07 |
| | A3-A3 | 0.33±0.05 | 0.35±0.06 | 0.34±0.08 | 0.38±0.05 |
| | A3-2/5 | 0.49±0.06 | 0.21±0.06 | 0.29±0.07 | 0.32±0.04 |
| | A3-4 | 0.31±0.06 | 0.13±0.06 | 0.10±0.02 | 0.22±0.03 |
| | A2/5-1 | 0.54±0.04 | 0.33±0.01 | 0.25±0.04 | 0.29±0.03 |
| | A2/5-3 | 0.54±0.07 | 0.31±0.01 | 0.29±0.08 | 0.29±0.04 |
| | A2/5-2/5 | 0.50±0.01 | 0.44±0.02 | 0.33±0.01 | 0.39±0.01 |
| | A2/5-4 | 0.52±0.09 | 0.24±0.01 | 0.10±0.01 | 0.45±0.02 |
| | A4-1 | 0.50±0.02 | 0.26±0.01 | 0.25±0.02 | 0.44±0.09 |
| | A4-3 | 0.37±0.01 | 0.24±0.01 | 0.12±0.09 | 0.36±0.08 |
| | A4-2/5 | 0.53±0.09 | 0.16±0.03 | 0.37±0.09 | 0.63±0.04 |
| | A4-4 | 0.52±0.03 | 0.52±0.01 | 0.27±0.04 | 0.29±0.01 |
| | Average | 0.42 | 0.27 | 0.26 | 0.37 |
| α-1,3-glucan ($A^b$) | A1-1 | 0.35±0.02 | 0.99±0.01 | 0.34±0.04 | 0.38±0.03 |
| | A1-3 | 0.68±0.01 | 0.11±0.06 | 0.41±0.01 | 0.39±0.07 |
| | A1-4 | 0.88±0.06 | 0.55±0.01 | 0.29±0.01 | 0.35±0.04 |
| | A3-A1 | 0.14±0.03 | 0.23±0.01 | 0.31±0.01 | 0.32±0.07 |
| | A3-3 | 0.40±0.06 | 0.37±0.06 | 0.34±0.08 | 0.38±0.05 |
| | A3-4 | 0.13±0.01 | 0.48±0.02 | 0.24±0.06 | 0.28±0.04 |
| | A4-1 | 0.39±0.04 | 0.31±0.06 | 0.21±0.01 | 0.43±0.05 |
| | A4-3 | 0.40±0.06 | 0.24±0.07 | 0.29±0.02 | 0.39±0.05 |
| | A4-4 | 0.49±0.01 | 0.42±0.01 | 0.32±0.08 | 0.43±0.01 |
| | Average | 0.43 | 0.41 | 0.31 | 0.37 |
| α-1,3-glucan ($A^c$) | A1-1 | 0.61±0.03 | 0.23±0.01 | 0.37±0.04 | 0.53±0.04 |
| | A1-3 | 0.94±0.08 | 0.57±0.07 | 0.21±0.02 | 0.61±0.07 |
| | A1-4 | 0.02±0.09 | 0.01±0.01 | 0.69±0.08 | 0.99±0.08 |
| | A3-A1 | 0.87±0.03 | 0.17±0.01 | 0.53±0.04 | 0.68±0.06 |
| | A3-3 | 0.53±0.01 | 0.58±0.03 | 0.14±0.02 | 0.42±0.05 |
| | A3-4 | 0.99±0.04 | 0.49±0.02 | 0.36±0.01 | 0.83±0.01 |
| | A4-1 | 0.43±0.01 | 0.34±0.01 | 0.37±0.09 | 0.34±0.03 |
| | A4-3 | 0.33±0.02 | 0.23±0.01 | 0.97±0.01 | 0.46±0.03 |
| | A4-4 | 0.52±0.02 | 0.33±0.02 | 0.36±0.08 | 0.44±0.06 |
| | Average | 0.58 | 0.33 | 0.45 | 0.59 |
| β-1,6-glucan (H) | H3-3 | 0.87±0.02 | 0.50±0.04 | 0.44±0.01 | 0.47±0.01 |
| | H3-5 | 0.97±0.02 | 0.63±0.03 | 0.44±0.01 | 0.51±0.02 |
| | H3-2 | 0.98±0.06 | 0.30±0.02 | 0.66±0.03 | 0.46±0.04 |
| | H5-3 | 0.47±0.01 | 0.65±0.01 | 0.36±0.09 | 0.33±0.01 |
| | H5-2 | 0.62±0.03 | 0.55±0.01 | 0.51±0.01 | 0.50±0.02 |
| | H5-5 | 0.64±0.03 | 0.74±0.02 | 0.78±0.04 | 0.50±0.03 |
| | H2-3 | 0.82±0.09 | 0.65±0.04 | 0.92±0.01 | 0.57±0.08 |

| | | | | | |
|---|---|---|---|---|---|
| | H2-5 | 0.98±0.05 | 0.48±0.02 | 0.94±0.05 | 0.89±0.05 |
| | H2-2 | 0.61±0.01 | 0.52±0.01 | 0.56±0.01 | 0.56±0.01 |
| | Average | 0.77 | 0.56 | 0.62 | 0.53 |
| Chitin (Ch) | Ch1-1 | 0.57±0.04 | 0.10±0.01 | 0.41±0.01 | 0.27±0.02 |
| | Ch1-3 | 0.93±0.06 | 0.12±0.02 | 0.54±0.03 | 0.68±0.02 |
| | Ch1-2 | 0.88±0.04 | 0.41±0.02 | 0.49±0.02 | 0.59±0.03 |
| | Ch3-1 | 0.79±0.02 | 0.54±0.03 | 0.45±0.02 | 0.72±0.01 |
| | Ch3-3 | 0.85±0.02 | 0.34±0.02 | 0.36±0.02 | 0.64±0.02 |
| | Ch3-2 | 0.64±0.04 | 0.53±0.08 | 0.91±0.02 | 0.62±0.05 |
| | Ch2-1 | 0.93±0.01 | 0.71±0.02 | 0.02±0.04 | 0.99±0.01 |
| | Ch2-3 | 0.64±0.07 | 0.53±0.03 | 0.68±0.02 | 0.91±0.06 |
| | Ch2-2 | 0.51±0.02 | 0.58±0.01 | 0.59±0.07 | 0.56±0.01 |
| | Average | 0.75 | 0.43 | 0.49 | 0.67 |
| Mannan (Mn[1,2,3]) | Mn1-1 | 0.32±0.05 | - | 0.35±0.05 | - |
| | Mn1-3 | 0.49±0.08 | - | 0.16±0.03 | - |
| | Mn1-5 | 0.65±0.06 | - | 0.52±0.09 | - |
| | Mn3-1 | 0.42±0.04 | - | 0.27±0.03 | - |
| | Mn3-3 | 0.56±0.02 | - | 0.3±0.04 | - |
| | Mn3-5 | 0.65±0.05 | - | 0.19±0.01 | - |
| | Mn5-1 | 0.15±0.01 | - | 0.29±0.01 | - |
| | Mn5-3 | 0.22±0.01 | - | 0.99±0.09 | - |
| | Mn5-5 | 0.01±0.01 | - | 0.56±0.04 | - |
| | Average | 0.38 | | 0.41 | |

**Supplementary Table 6. $^1$H-T$_{1\rho}$ and $^{13}$C-T$_1$ relaxation times of polysaccharides in cell walls.** Data is shown for the *C. neoformans* samples. The average values for each molecule in each sample are highlighted in bold. The data were measured using 1D $^{13}$C relaxation experiments. The data are fit using single exponential equations: $I(t) = e^{-t/T_1}$. Error bars are standard deviations of the fit parameters.

| Polysaccharide | Chemical shift | H99 | | *pka1* | | JEC20 | | Cap70 | |
|---|---|---|---|---|---|---|---|---|---|
| | | $^1$H-T$_{1\rho}$ (ms) | $^{13}$C-T$_1$ (s) | $^1$H-T$_{1\rho}$ (ms) | $^{13}$C-T$_1$ (s) | $^1$H-T$_{1\rho}$ (ms) | $^{13}$C-T$_1$ (s) | $^1$H-T$_{1\rho}$ (ms) | $^{13}$C-T$_1$ (s) |
| α-1,3-glucan (a) | 101.5 | 11.4±1.2 | 3.6±0.2 | 13.6±1.0 | 3.4±0.1 | 18.6±1.3 | 3.8±0.2 | 14.0±1.1 | 4.8±0.4 |
| | 85.0 | 12.5±1.0 | 3.5±0.2 | 15.0±1.4 | 3.8±0.1 | 22.0±1.4 | 4.3±0.1 | 16.0±1.0 | 4.8±0.4 |
| | 71.9 | 8.5±0.9 | 2.8±0.3 | 9.8±0.9 | 2.5±0.2 | 12.4±1.0 | 3.0±0.2 | 10.8±0.9 | 3.3±0.3 |
| | 69.6 | 6.5±0.8 | 2.1±0.3 | 8.5±1.0 | 1.3±0.2 | 13.0±1.2 | 2.6±0.3 | 9.6±1.0 | 2.6±0.4 |
| | Average | 9.7 | 3.0 | 11.7 | 2.7 | 16.4 | 3.4 | 12.6 | 3.9 |
| Chitin | 104.6 | 4.9±0.6 | 1.3±0.1 | 6.9±1.1 | 2.5±0.5 | 4.7±0.8 | 1.5±0.3 | 7.9±1.2 | 2.2±0.4 |
| | 83.4 | 6.4±0.7 | 2.0±0.2 | 9.5±1.3 | 2.5±0.1 | 8.3±1.1 | 2.2±0.1 | 10.2±0.8 | 2.8±0.3 |
| | 55.9 | 3.8±0.5 | 0.9±0.1 | 7.6±0.8 | 1.4±0.09 | 8.0±0.9 | 1.0±0.1 | 7.3±1.0 | 1.4±0.2 |
| | Average | 5.1 | 1.4 | 8.0 | 2.1 | 7.0 | 1.6 | 8.5 | 2.1 |
| β-1,6-glucan | 103.9 | 4.2±0.4 | 1.1±0.1 | 5.9±0.7 | 0.7±0.08 | 6.3±0.6 | 1.4±0.2 | 5.0±0.6 | 1.0±0.2 |
| | 76.4 | 3.2±0.2 | 0.8±0.07 | 4.4±0.5 | 0.6±0.06 | 4.2±0.4 | 0.9±0.1 | 4.1±0.3 | 0.8±0.1 |
| | 75.6 | 3.4±0.3 | 0.9±0.09 | 4.7±0.5 | 0.8±0.08 | 4.3±0.5 | 1.4±0.1 | 4.5±0.4 | 1.1±0.1 |
| | 74.0 | 4.0±0.3 | 1.0±0.1 | 5.8±0.6 | 0.9±0.1 | 6.6±0.6 | 1.4±0.2 | 5.5±0.5 | 1.2±1.4 |
| | 70.6 | 7.0±0.8 | 2.0±0.2 | 7.9±0.8 | 1.3±0.2 | 13.0±0.9 | 2.4±0.2 | 9.2±0.9 | 2.4±0.3 |
| | Average | 4.4 | 1.2 | 5.8 | 0.9 | 6.8 | 1.5 | 5.6 | 1.3 |

**Supplementary Table 7. $^{1}$H and $^{13}$C chemical shifts of *C. neoformans* polysaccharides from $^{1}$H-detected experiments.** For each carbon site, the $^{13}$C and $^{1}$H chemical shifts are shown in the top and bottom rows, respectively. The referencing scale is TMS scale for $^{13}$C, and DSS for $^{1}$H. Underline: sites with ambiguity due to spectral overlap.

| Carbohydrates | forms | C1/H1 | C2/H2 | C3/H3 | C4/H4 | C5/H5 | C6/H6 |
|---|---|---|---|---|---|---|---|
| Rigid molecules | | | | | | | |
| α-1,3-glucan (A) | a | 101.6<br>5.5 | 71.2<br>4.2 | 85.5<br>3.5 | 71.2<br>4.2 | 71.2<br>4.2 | 61.2<br>4.4 |
| | b | 101.4<br>4.8 | 71.5<br>3.6 | 85.0<br>3.8 | 71.5<br>3.6 | 71.5<br>3.6 | 61.5<br>3.9 |
| | e | 101.2<br>6.1 | 71.5<br>3.2 | 85.0<br>4.4 | 71.5<br>3.2 | 71.5<br>3.2 | 61.2<br>3.4 |
| β-1,6-glucan (H) | | 104<br>5.4-3.7 | 74.2<br>4.7-3.1 | 76.8<br>4.7-3.1 | 70.8<br>4.7-3.0 | 75.8<br>4.7-2.6 | 69.8<br>4.7-2.8 |

**Supplementary Table 8. $^{13}C$ chemical shifts of *C. neoformans* melanin from $^{1}H$-detected experiments.** For each aromatic carbon site, the $^{13}C$ chemical shifts are shown in ppm.

| Melanin fragments | C4 | C5 | C6 | C7 | C8 | C9 |
|---|---|---|---|---|---|---|
| A | 110.0-125.0 | 135.0 | 148.0 | 137.0 | 145.0 | 140.0-144.0 |
| B | / | 145.0 | 140.0-144.0 | / | 125.0-130.0 | 110.0-125.0 |
| C | 125.0-130.0 | 153.0 | 175.0-178.0 | / | 168.5 | 140.0-144.0 |
| D | 110.0-125.0 | 137.0 | 140.0-144.0 | / | 125.0-130.0 | 140.0-144.0 |

**Supplementary Table 9. $^{13}$C chemical shifts of *C. neoformans* melanin from literature.** For each group of aromatic carbons, the $^{13}$C chemical shifts are shown in ppm.

| Observed shifts (ppm) | Chemical grouping | Literature referencing |
|---|---|---|
| 110-118 | Aromatic -CH- | Vecchia *et al.* 2013[10]<br>Adhyaru *et al.* 2003[11]<br>Johnson *et al.* 2013[12] |
| 125-130 | -CH=CH-, indole or alkene | Vecchia *et al.* 2013[10]<br>Adhyaru *et al.* 2003[11]<br>Johnson *et al.* 2013[12] |
| 144-156 | Aromatic -CH=CHCO-, aromatic -C- | Vecchia *et al.* 2013[10]<br>Johnson *et al.* 2013[12]<br>Subhasish *et al.* 2014[13] |
| 157-165 | Aromatic -C- | Vecchia *et al.* 2013[10]<br>Johnson *et al.* 2013[12] |
| 170-180 | -COO-, -CONH | Vecchia *et al.* 2013[10]<br>Subhasish *et al.* 2014[13] |

**Supplementary Table 10. $^{13}$C Solid-state NMR experimental parameters for fungal cell wall characterization.** T = sample temperature; $B_0$ = magnetic field; $\nu_{MAS}$ = MAS frequency; ns = number of scans; $d_1$ = recycle delay between scans; $t_{1, max}$ = maximum $t_1$ evolution time (for indirect dimension); $t_{1, inc}$ = increment for $t_1$ (for indirect dimension) evolution time; $\tau_{dw}$ = dwell time during direct FID acquisition; $\tau_{acq}$ = maximum acquisition time during direct FID detection; $\tau_{XY}$ = cross-polarization contact time during CP from channel X to channel Y; $\nu_{1H, dec}$ = dipolar decoupling field strength**.** Spin diffusion (SD).

| Experiment | NMR Parameters | | | | | | | | | | | | | | | Samples |
|---|---|---|---|---|---|---|---|---|---|---|---|---|---|---|---|---|
| | T (K) | $B_0$ (T) | $\nu_{MAS}$ (kHz) | ns | d1 (s) | $t_{1, max}$ (ms) | $t_{1, inc}$ (μs) | $\tau_{dw}$ (μs) | $\tau_{acq}$ (ms) | $\tau_{HC}$ (ms) | $\tau_{HN}$ (ms) | $\tau_{NC}$ (ms) | $\tau_{SD}$ (ms) | $\tau_{mix}$ (ms) | $\nu_{1H\ dec}$ (k Hz) | |
| Identification and quantification of polysaccharides | | | | | | | | | | | | | | | | H99, JEC20, *pka1*, Cap70 |
| 1D $^{13}$C CP | 298 | 18.8 | 15 | 2048 | 2 | | | 5 | 18 | 1 | | | | | 83 | |
| 1D $^{13}$C DP | 298 | 18.8 | 15 | 512 | 2 or 35 | | | 5 | 18 | | | | | | 83 | |
| 1D $^{13}$C refocused INEPT | 298 | 18.8 | 15 | 1024 | 3.5 | | | 5 | 16 | | | | | 1.7 $\tau_J$ | 83 | |
| 2D $^{13}$C-$^{13}$C with CORD mixing | 298 | 18.8 | 15 | 32 | 2 | 7.5 | 25 | 5 | 14 | 0.5 | | | | 53 $\tau_{CORD}$ | 83 | |
| 2D $^{13}$C-$^{13}$C refocused CP J-INADEQUATE | 298 | 18.8 | 15 | 16 | 2 | 7.5 | 22 | 5 | 14 | | | | | | 83 | |
| 2D $^{13}$C-$^{13}$C refocused DP J-INADEQUATE | 298 | 18.8 | 15 | 16 | 2 | 7.5 | 22 | 5 | 14 | | | | | | 83 | |
| Estimation of site-specific hydration of polysaccharides | | | | | | | | | | | | | | | | |
| 2D $^{13}$C-$^{13}$C water-edited | 280 | 9.4 | 15 | 64 | 2 | 5.5 | 50 | 8 | 16 | 1 | | | 0, 4 | 50 $\tau_{PDSD}$ | 71 | |
| Dynamics of polysaccharides | | | | | | | | | | | | | | | | |
| 1D $^{13}$C-$T_1$ | 298 | 9.4 | 15 | 512 | 2 | | | 8 | 16 | 1 | | | | | 71 | |
| 1D $^1$H-$T_{1\rho}$ | 298 | 9.4 | 15 | 512 | 2 | | | 8 | 16 | 1 | | | | | 71 | |
| 1D gated CP | 298 | 14.1 | 14 | 1024 | 2 | | | 10 | 16 | 5 | | | | | 83 | H99 |

**Supplementary Table 11. Parameters used for proton detection experiments.** The CP based proton detection experiments were performed on 600 MHz (14.1 T) spectrometer with the MAS frequency of 60 kHz.

| Expt. | Temp. (K) | CP (μs) | | D1 | NS | td2 | td1 | td3 | aq2 (ms) | aq1 (ms) | aq3 (ms) | Water suppression | TOCSY(WALTZ-16) (ms) | RFDR Mixing (ms) | Sample |
|---|---|---|---|---|---|---|---|---|---|---|---|---|---|---|---|
| | | $t_{cp1}$ | $t_{cp2}$ | | | | | | | | | | | | |
| 2D hCH | 304 | 1000 (HC-CP) | 100 (CH-CP) | 3 | 32 | 1764 ($^{1}H$) | 320 ($^{13}C$) | - | 14.9 | 5.3 | - | MISSISSIPI (total duration) 100 ms (rf 15.2 kHz) | - | - | Melanized *C. neoformans* H99 |
| 2D hChH (RFDR) | 304 | 1000 (HC-CP) | 500 (CH-CP) | 2 | 512 | 1600 ($^{1}H$) | 192 ($^{13}C$) | - | 13.6 | 2.39 | - | | - | 0.8 | Melanized *C. neoformans* H99 |
| 3D hCCH TOCSY (WALTZ-16) | 304 | 1000 | 100 | 2 | 8 | 1764 | 128 | 128 | 14.9 | 2.13 | 2.13 | MISSISSIPI (total duration) 100 ms (rf 15.2 kHz | 15 ms (rf 21.4 kHz) | - | Melanized *C. neoformans* H99 |

### Supplementary Reference

1 Chakraborty, A. *et al.* A molecular vision of fungal cell wall organization by functional genomics and solid-state NMR. *Nat. Commun.* **12**, 6346 (2021).
2 Lowman, D. *et al.* New Insights into the Structure of (1→3,1→6)-β-D-Glucan Side Chains in the *Candida glabrata* Cell Wall. *PLoS ONE* **6**, e27614 (2011).
3 Shim *et al.* Antitumor effect of soluble β-1,3-glucan from Agrobacterium sp r259 KCTC 1019. *J. Microbiol. Biotechnol.* **17**, 1513-1520 (2007).
4 Fairweather, J. K., Him, J. L. K., Heux, L., Driguez, H. & Bulone, V. Structural characterization by $^{13}$C-NMR spectroscopy of products synthesized in vitro by polysaccharide synthases using $^{13}$C-enriched glycosyl donors: application to a UDP-glucose:(1→ 3)-b-d-glucan synthase from blackberry (Rubus fruticosus). *Glycobiology* **14**, 775-781 (2004).
5 Saitô, H., Ohki, T. & Sasaki, T. A 13C-nuclear magnetic resonance study of polysaccharide gels. Molecular architecture in the gels consisting of fungal, branched (1→ 3)-b-D-glucans (lentinan and schizophyllan) as manifested by conformational changes induced by sodium hydroxide. *Carbohydr. Res.* **74**, 227-240 (1979).
6 Kang, X. *et al.* Molecular architecture of fungal cell walls revealed by solid-state NMR. *Nat. Commun.* **9**, 2747 (2018).
7 Fernando, L. D. *et al.* Structural Polymorphism of Chitin and Chitosan in Fungal Cell Walls From Solid-State NMR and Principal Component Analysis. *Front. Mol. Biosci.* **8**, 727053 (2021).
8 Bacon, B., Cherniak, R., KwonChung, K. & Jacobson, E. Structure of the O-deacetylated glucuronoxylomannan from *Cryptococcus neoformans* Cap70 as determined by 2D NMR spectroscopy. *Carbohydr. Res.* **283**, 95-110 (1996).
9 Moyrand, F., Klaproth, B., Himmelreich, U., Dromer, F. & Janbon, G. Isolation and characterization of capsule structure mutant strains of *Cryptococcus neoformans*. *Mol. Microbiol.* **45**, 837-849 (2002).
10 Della Vecchia, N. *et al.* Building-Block Diversity in Polydopamine Underpins a Multifunctional Eumelanin-Type Platform Tunable Through a Quinone Control Point. *Adv. Funct. Mater.* **23**, 1331-1340 (2013).
11 Adhyaru, B., Akhmedov, N., Katritzky, A. & Bowers, C. Solid-state cross-polarization magic angle spinning 13C and 15N NMR characterization of Sepia melanin, Sepia melanin free acid and Human hair melanin in comparison with several model compounds. *Magn. Reson. Chem.* **41**, 466-474 (2003).
12 Johnson, R. *et al.* Spectrally edited 2D 13C-13C NMR spectra without diagonal ridge for characterizing 13C-enriched low-temperature carbon materials. *J. Magn. Reson.* **234**, 112-124 (2013).
13 Chatterjee, S. *et al.* Demonstration of a common indole-based aromatic core in natural and synthetic eumelanins by solid-state NMR. *Org. Biomol. Chem.* **12**, 6730-6736 (2014).
